# Uterine ERBB3 signaling is critical to cultivate stromal environment for functional gland branching

**DOI:** 10.1101/2025.10.16.682855

**Authors:** Bo Li, Amanda Dewar, Sudhansu K. Dey, Xiaofei Sun

## Abstract

Preimplantation uterine gland differentiation, involving glandular branching, is a prerequisite for secretion of leukemia inhibitory factor (LIF), thereby establishing uterine receptivity to embryo implantation. This differentiation process is disrupted by the absence of FOXA2, a transcription factor specifically present in uterine glands. It remains unclear whether preimplantation glandular differentiation is regulated exclusively by gland-intrinsic mechanisms or is influenced by the surrounding microenvironment through mesenchymal-epithelial interactions. We show here that uterine deletion of *Erbb3* leads to impaired gland branching and FOXA2 expression, as well as reduced female fertility. In contrast, females with uterine epithelial-specific deletion of *Erbb3* show normal fertility, suggesting a critical role of stromal ERBB3 signaling. A transcriptomic profiling of day 3 stromal cells identified *Igf1* as a candidate paracrine factor mediating the stromal-epithelial interaction. Thus, the uterine deletion of *Igf1r* leads to a reduced gland branching phenotype similar to that observed in *Erbb3^d/d^* uteri. These results demonstrate that ERBB3 regulates glandular branching and embryo implantation via the ERBB3-IGF1-IGF1R signaling pathway.

## Introduction

Pregnancy is a tightly synchronized process, comprising implantation, decidualization, placentation and parturition ^1, 2^. Successful embryo implantation requires cross-talk between the implantation-competent blastocyst and the receptive uterus ^3, 4^. The uterus is composed of the endometrium and myometrium, with the endometrium consisting of stromal, glandular, and luminal epithelial compartments. Uterine glands play a critical role in establishing uterine receptivity ^5^. In both mice and humans, these glands synthesize and secrete factors required for uterine receptivity, embryo implantation, and subsequent embryo development ^6, 7^. In mice, leukemia inhibitory factor (LIF) is a key gland-derived cytokine secreted immediately before implantation on day 4 of pregnancy (Day 1 = morning of finding vaginal plug). LIF can substitute for nidatory estrogen to induce implantation in both normal and delayed implantation models ^8, 9^, and loss of maternal LIF results in complete implantation failure ^10^.

In both mouse and human uteri, the transcription factor Forkhead box A2 (FOXA2) is specifically expressed in glandular epithelial cells ^11, 12^. Genetic ablation of *Foxa2* in the whole uterus or uterine epithelium results in implantation failure due to impaired LIF production ^5, 13, 14^, underscoring a critical role for FOXA2 in glandular function and implantation success. Recent advances in tissue clearing and three-dimensional imaging have enabled detailed visualization of the uterine gland architecture ^15, 16^. Using these approaches, we previously demonstrated that uterine glands undergo a FOXA2-dependent differentiation program characterized by extensive branching during the preimplantation period, a process that prepares glands for estrogen responsiveness and subsequent LIF production ^17^. This defective gland branching phenotype was observed in mice with epithelial deletion of the estrogen receptor *Esr1* using *Pax2*-Cre or *Wnt7a*-Cre ^18^. To date, however, regulatory factors governing gland branching have been identified almost exclusively within glandular epithelial cells. Given that gland branching occurs within the stromal compartment, an important unanswered question is whether surrounding stromal cells contribute to this morphogenetic process. Moreover, although FOXA2 serves as a putative marker of uterine glands, glandular epithelium originates from FOXA2-negative luminal epithelium, raising the possibility that FOXA2 expression is not solely cell-autonomous but is instead influenced by cues from the neighboring stromal microenvironment.

Epidermal growth factors (EGFs) and their receptors are critical for pregnancy ^19, 20, 21^. EGF signaling is mediated through the ERBB family of receptor tyrosine kinases (ERBB1-4), which function as homo- or heterodimers ^22, 23^. ERBB3 lacks intrinsic kinase activity distinguishing it from other ERBB receptors. Previous studies have shown that *Erbb3* is expressed in both epithelial and stromal cells in the peri-implantation mouse uterus ^24^. Our previous study identified ERBB2 and ERBB3 as major receptors for heparin-binding EGF (HB-EGF) in early pregnancy ^20^. In the study, we generated females with uterine-specific deletion of *Erbb3* (*Erbb3^f/f^ Pgr^Cre/+^*, hereafter *Erbb3^d/d^*). Notably, around 70% of vaginal plug-positive *Erbb3^d/d^* females failed to produce live offspring. A closer examination revealed a marked reduction in uterine gland branching in *Erbb3^d/d^* uteri, leading us to hypothesize that uterine ERBB3 signaling regulates gland branching and function, thereby influencing female fertility and pregnancy success.

In this study, we demonstrate that *Erbb3^d/d^* females exhibit implantation failure associated with reduced LIF secretion. Uterine deletion of *Erbb3* disrupts preimplantation gland branching and impairs glandular function required for the establishment of uterine receptivity. Interestingly, FOXA2 expression is markedly diminished in uterine glands of *Erbb3^d/d^* females, suggesting that ERBB3 signaling is essential for maintaining glandular cell identity. Transcriptomic analysis of day 3 uteri further reveals a significant reduction in stromal *Igf1* expression in *Erbb3^d/d^* mice, and uterine deletion of *Igf1r* recapitulates the glandular branching defects observed in *Erbb3^d/d^*uteri, suggesting that stromal IGF1 signaling acts as a key downstream mediator of ERBB3-dependent regulation of uterine gland development and function.

## Results

### Simultaneous epithelial and stromal deletion of *Erbb3* compromises embryo implantation

In our previous study of uterine HB-EGF ^20^, we generated *Erbb3^d/d^* female mice with deletion of *Erbb3* in uterine stromal and epithelial cells by crossing *Erbb3^flox/flox^* mice with *Pgr^Cre/+^* mice, and females with uterine epithelial-specific deletion of *Erbb3* by crossing *Erbb3^flox/flox^* mice with *Ltf^Cre/+^* mice (*Erbb3^epi/epi^*). Deletion efficiency was confirmed by Western blot analysis using day 4 *Erbb3^d/d^* uterine tissues (Supplementary Fig. 1a, b). Since *Erbb3* is expressed in epithelial and stromal cells in preimplantation uteri ^24^, we further assessed compartment-specific deletion efficiency by isolating primary stromal and epithelial cells from *Erbb3^f/f^* and *Erbb3^d/d^* uteri. qPCR analyses demonstrated robust depletion of *Erbb3* in both compartments of *Erbb3^d/d^*uteri (Supplementary Fig. 1c-e).

Consistent with our previous findings, approximately 70% of plug-positive *Erbb3^d/d^* females failed to deliver offspring ^20^, whereas *Erbb3^epi/epi^* females have pregnancy rates and litter sizes comparable to those of *Erbb3^f/f^*females (Fig. 1a and Supplementary Table 1). We next investigated the developmental onset of this infertility. While *Erbb3^f/f^*and *Erbb3^epi/epi^* females exhibited normal embryo attachment, 57% (8/14) of *Erbb3^d/d^* females showed implantation failure on day 5 of pregnancy (Fig. 1b, c). Recovery of blastocysts from these females indicates that the implantation failure is due to a maternal defect. This divergent reproductive outcome suggests that although epithelial *Erbb3* is dispensable for implantation, the combined loss of *Erbb3* in both epithelial and stromal compartments markedly compromises pregnancy success. Taken together, these findings implicate stromal *Erbb3* as a critical regulator of female fertility.

**Fig. 1.**
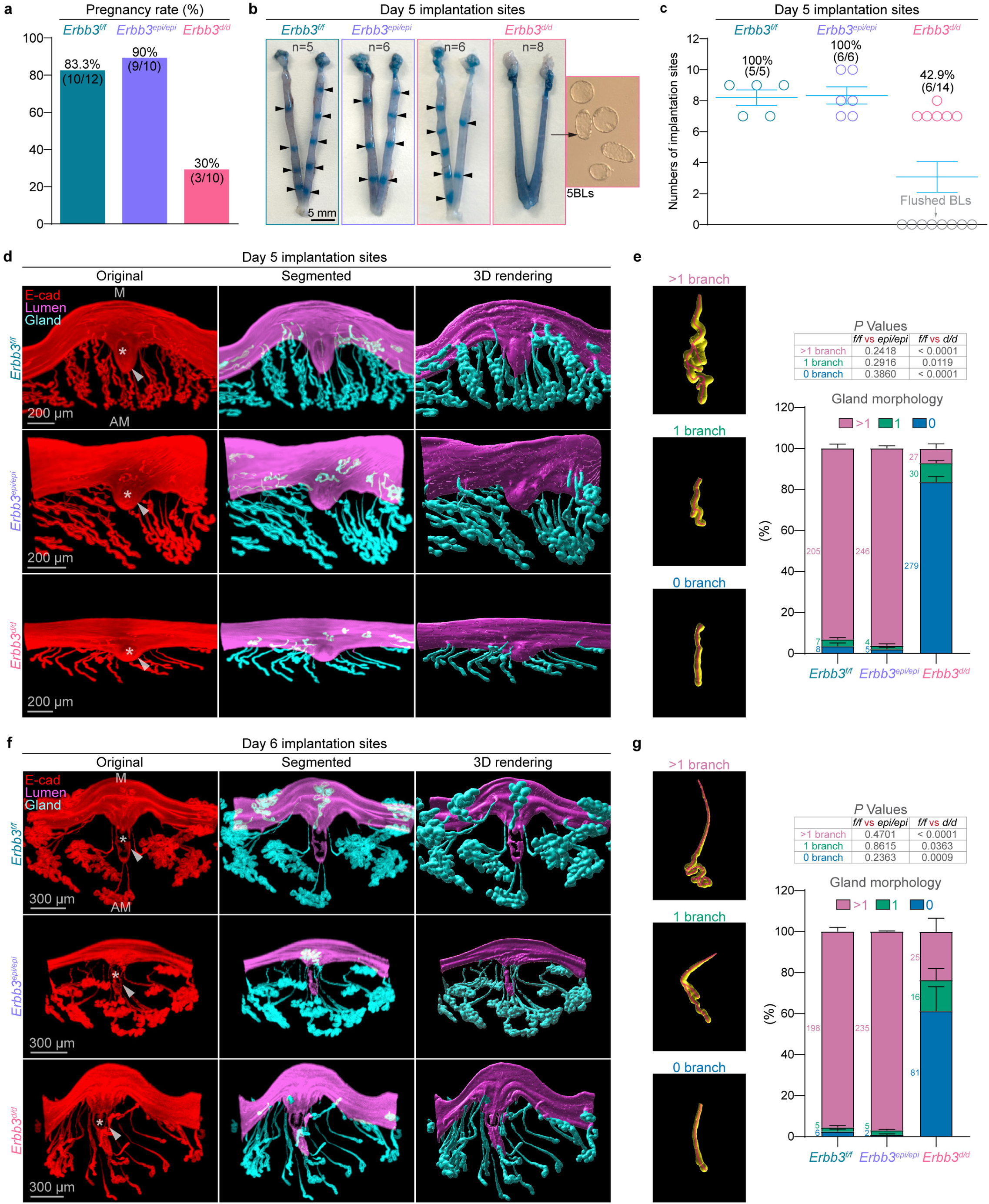
Simultaneous epithelial and stromal deletion of *Erbb3* compromises embryo implantation. **a** Pregnancy rates of *Erbb3^f/f^* (n = 12), *Erbb3^epi/epi^* (n = 10), and *Erbb3^d/d^* (n = 10) mice. Pregnancy rate is defined as the percentage of vaginal plug-positive mice that successfully delivered pups. Source data are provided as a Source Data file. **b** Representative uteri from *Erbb3^f/f^*, *Erbb3^epi/epi^*, and *Erbb3^d/d^* females after blue dye injection on day 5. Blastocysts were recovered by flushing uterine horns from the *Erbb3^d/d^*uteri without blue bands. BL, blastocysts. Arrowheads indicate implantation sites. **c** Numbers of day 5 implantation sites from *Erbb3^f/f^* (n = 5), *Erbb3^epi/epi^* (n = 6), and *Erbb3^d/d^* (n = 14) mice. Each circle represents an individual mouse. Source data are provided as a Source Data file. **d** 3D imaging of day 5 implantation sites in *Erbb3^f/f^* (n = 4), *Erbb3^epi/epi^* (n = 5), and *Erbb3^d/d^* (n = 8) females. Images of E-cadherin immunostaining, segmented, and 3D rendering of day 5 implantation sites in each genotype. Scale bar, 200 μm. Asterisks indicate the location of blastocysts. Arrowheads indicate the implantation chamber (crypt). M mesometrial pole, AM antimesometrial pole. **e** Quantification of gland branches at day 5 implantation sites from *Erbb3^f/f^* (n = 4), *Erbb3^epi/epi^* (n = 5), and *Erbb3^d/d^*(n = 8) mice. Data are represented as mean ± SEM. *P* values were determined by a two-tailed Student’s *t* test. Source data are provided as a Source Data file. **f** 3D imaging of day 6 implantation sites in *Erbb3^f/f^*(n = 5), *Erbb3^epi/epi^* (n = 6), and *Erbb3^d/d^*(n = 4) females. Images of E-cadherin immunostaining, segmented, and 3D rendering of day 6 implantation sites in each genotype. Scale bars, 300 μm. Asterisks indicate the location of blastocysts. Arrowheads indicate the implantation chamber (crypt). M mesometrial pole, AM antimesometrial pole. **g** Quantification of gland branches at day 6 implantation sites from *Erbb3^f/f^* (n = 5), *Erbb3^epi/epi^* (n = 6), and *Erbb3^d/d^* (n = 4) females. Data are represented as mean ± SEM. *P* values were determined by a two-tailed Student’s *t* test. Source data are provided as a Source Data file.

### Uterine deletion of *Erbb3* impairs gland branching

On day 5 of pregnancy, blastocysts attach to the luminal epithelium and nidate into a spear-shaped implantation chamber (crypt) that is connected with adjacent uterine glands ^16^. Using 3D imaging, we assessed the chamber morphology of *Erbb3^f/f^*, *Erbb3^epi/epi^* and *Erbb3^d/d^* implantation sites on days 5 and 6 of pregnancy. On day 5, *Erbb3^d/d^* implantation sites exhibited shallower crypt invaginations compared with those of *Erbb3^f/f^* and *Erbb3^epi/epi^* implantation sites (Fig. 1d). Interestingly, while glands in *Erbb3^f/f^* and *Erbb3^epi/epi^* uteri displayed multiple branches on day 5 of pregnancy, most *Erbb3^d/d^* uterine glands retained a single, elongated structure lacking typical branching (Fig. 1d). Additional implantation chamber images are shown in Supplementary Fig. 2. Glands were categorized as unbranched (stem only, 0 branches), containing one or more than one branch (Fig. 1e). Quantitative analysis revealed that the proportion of *Erbb3^d/d^* glands with one or more branches was significantly reduced compared with control glands. Development of the *Erbb3^d/d^* crypts rapidly progressed, and by day 6 of pregnancy, no overt defects in crypt formation were observed across different genotypes (Fig. 1f). At this stage,*Erbb3^f/f^* and *Erbb3^epi/epi^* crypts showed continued extension of glandular stems accompanied by coiled branches that were displaced away from the implantation chambers. In contrast, most *Erbb3^d/d^* uterine glands remained as elongated main ducts with minimal or no branching (Fig. 1f, g and Supplementary Fig. 3). These results suggest that uterine gland branching is defective in *Erbb3^d/d^* implantation sites.

### Uterine deletion of *Erbb3* reduces LIF secretion

To investigate the cause of impaired implantation in *Erbb3^d/d^* females, we examined ovarian hormone levels and their putative receptor expression for uterine receptivity. The ovarian hormones, 17β-Estradiol (E_2_) and progesterone (P_4_), play key roles in pregnancy ^25, 26^, and exert their functions primarily through estrogen receptor α (ESR1) and progesterone receptors (PR), respectively ^27^. Serum levels of E_2_ and P_4_, as well as uterine expression of their respective receptors, were comparable between *Erbb3^f/f^* and *Erbb3^d/d^* females on day 4 of pregnancy (Fig. 2a-c). *Erbb3^d/d^*uteri showed expression levels of *Msx1*, a well-established marker of uterine receptivity ^28^, comparable to those of control on day 4 of pregnancy (Fig. 2d), despite a reduced number of uterine glands observed in *Erbb3^d/d^* sections. Under normal conditions on day 4 of pregnancy, luminal and glandular epithelial cells exit the cell cycle, whereas stromal cells remain highly proliferative ^29^. While *Erbb3^f/f^* uteri displayed this expected pattern, *Erbb3^d/d^* uteri exhibited aberrant Ki67 staining in luminal and glandular epithelia. In contrast, no significant differences in stromal Ki67 staining were detected between *Erbb3^f/f^*and *Erbb3^d/d^* uteri (Fig. 2e).

**Fig. 2.**
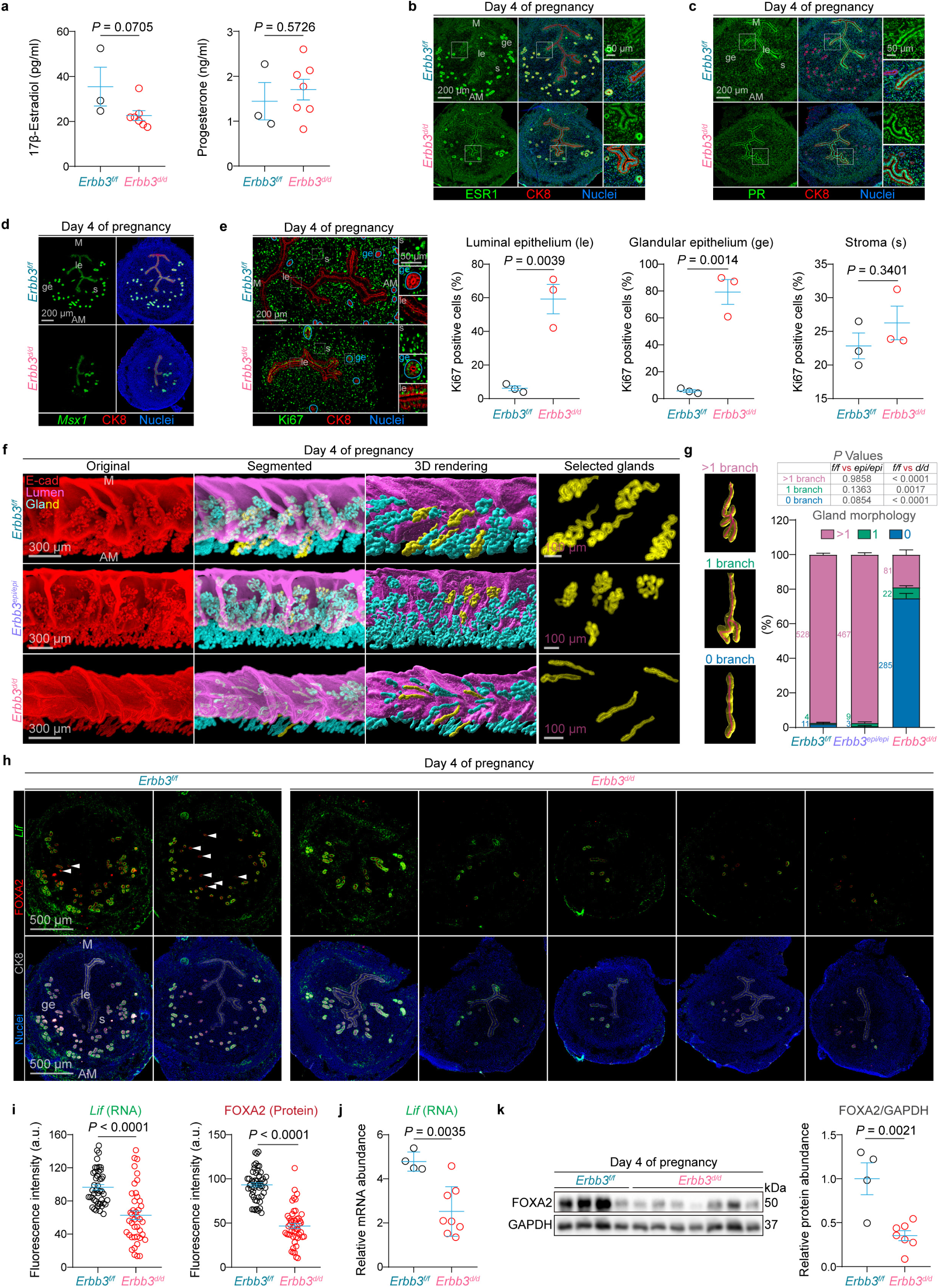
*Erbb3* deletion reduces *Lif* mRNA and FOXA2 protein expression. **a** Serum levels of 17β-Estradiol and progesterone in *Erbb3^f/f^* (n = 3) and *Erbb3^d/d^* (n = 7) females. Data are represented as mean ± SEM. *P* values were determined by a two-tailed Student’s *t* test. Source data are provided as a Source Data file. **b, c** Immunofluorescence staining for ESR1 and PR co-stain with CK8 on day 4 *Erbb3^f/f^* (n = 3) and *Erbb3^d/d^* (n = 3) mouse uteri. Scale bars, 200 μm. **d** FISH of *Msx1* and immunofluorescence of CK8 on day 4 *Erbb3^f/f^* (n = 3) and *Erbb3^d/d^* (n = 3) mouse uteri. Scale bar, 200 μm. **e** Immunofluorescence staining for Ki67 and CK8 on day 4 uterine sections from *Erbb3^f/f^* (n = 3) and *Erbb3^d/d^* (n = 3) mice, with quantification of Ki67-positive cells (%) in the luminal epithelium, glandular epithelium, and stroma. Scale bars, 200 μm (main) and 50 μm (inset). le, luminal epithelium; ge, glandular epithelium; s, stroma; M, mesometrial pole; AM, antimesometrial pole. Data are represented as mean ± SEM. *P* values were determined by a two-tailed Student’s *t* test. Source data are provided as a Source Data file. **f** 3D imaging of day 4 mouse uteri in *Erbb3^f/f^* (n = 5), *Erbb3^epi/epi^* (n = 4), and *Erbb3^d/d^* (n = 7) females. Original (E-cadherin), segmented, and 3D rendering of day 4 uteri. Scale bars, 300 μm. Selected glands from *Erbb3^f/f^*and *Erbb3^d/d^* uteri are displayed from the right panel. Scale bars, 100 μm. **g** Quantification of gland branches in *Erbb3^f/f^* (n = 5), *Erbb3^epi/epi^* (n = 4), and *Erbb3^d/d^*(n = 7) females on day 4 of pregnancy. Data are represented as mean ± SEM. *P* values were determined by a two-tailed Student’s *t* test. Source data are provided as a Source Data file. **h** Representative image of FISH for *Lif* and immunofluorescence for FOXA2 and CK8 of day 4 uteri from pregnant *Erbb3^f/f^* (n = 3) and *Erbb3^d/d^* (n = 5) mice. Arrowheads indicate *Lif*-negative glands. Scale bars, 500 µm. le, luminal epithelium; ge, glandular epithelium; s, stroma; M, mesometrial pole; AM, antimesometrial pole. **i** Quantification of *Lif* RNA and FOXA2 protein fluorescence intensities (a.u.) in uterine glands from *Erbb3^f/f^* (n = 3) and *Erbb3^d/d^* (n = 5) mice on day 4 of pregnancy. Fluorescence intensity (a.u.) was quantified following the method described in the Methods section. Data are represented as mean ± SEM. *P* values were determined by a two-tailed Student’s t test. Source data are provided as a Source Data file. **j** Relative mRNA level of *Lif* by qPCR in *Erbb3^f/f^* (n = 4) and *Erbb3^d/d^* (n = 8) uteri on day 4 of pregnancy at 1100 h. Data are presented as mean ± SEM. *P* values were determined by a two-tailed Student’s *t* test. Source data are provided as a Source Data file. **k** Western blot analysis of FOXA2 protein levels in *Erbb3^f/f^* (n = 4) and *Erbb3^d/d^* (n = 7) uteri on day 4 of pregnancy. Data are presented as mean ± SEM. *P* values were determined by a two-tailed Student’s *t*-test. Source data are provided as a Source Data file.

Uterine glands undergo progressive branching morphogenesis and functional differentiation as the uterus acquires receptivity prior to implantation ^17^. This differentiation is critical for the production of LIF in response to preimplantation E_2_ secretion on the morning of day 4 of pregnancy ^10^. Our observations of defective *Erbb3^d/d^* glandular morphology on day 5 prompted us to examine uterine gland architecture on day 4 using 3D imaging. The majority of *Erbb3^f/f^* and *Erbb3^epi/epi^*uterine glands exhibited extensive branching, whereas *Erbb3^d/d^*glands had significantly fewer glands with more than one branch, indicating defective glandular differentiation and potential functional defects (Fig. 2f, g and Supplementary Fig. 4). Indeed, *Erbb3^d/d^* uteri displayed significantly reduced *Lif* expression with some inter-individual variability (Fig. 2h, i). Among the five *Erbb3^d/d^* uteri examined, one exhibited *Lif* expression comparable to that of control uteri, whereas the remaining samples showed a pronounced reduction in glandular *Lif* expression. This decrease in *Lif* production was further quantitatively validated by qPCR analysis using RNA isolated from whole *Erbb3^f/f^* and *Erbb3^d/d^* uteri on day 4 of pregnancy (Fig. 2j). As introduced above, FOXA2 plays a key role in preimplantation gland differentiation and LIF production. We therefore examined FOXA2 expression by immunostaining. In *Erbb3^f/f^* uteri, all glands were marked by FOXA2 staining, while *Lif* signals were detected in glandular epithelial cells located away from the uterine lumen.

In contrast, FOXA2 signals were markedly reduced in *Erbb3^d/d^* uteri (Fig. 2h, i). A quantitative Western blot analysis confirmed a significant reduction in FOXA2 protein levels in *Erbb3^d/d^* uteri (Fig. 2k).

Together, these results demonstrate that *Erbb3^d/d^* uteri have defective uterine gland branching and impaired function. The marked reduction in FOXA2 expression within *Erbb3^d/d^* glands suggests that ERBB3 signaling plays a key role in regulating FOXA2-dependent gland identity and differentiation. Moreover, the wide inter-individual variability in *Lif* and FOXA2 expression observed in *Erbb3^d/d^* uteri likely underlies the partial penetrance of the reproductive phenotype, providing an explanation for why approximately 30% of *Erbb3^d/d^* females are able to carry pregnancies to term.

### Postnatal 6-week-old *Erbb3^d/d^* females show normal uterine gland morphology and FOXA2 expression

We next asked whether the glandular defects observed in adult *Erbb3^d/d^* females arise from abnormalities during early postnatal uterine gland development. An important caveat when comparing the *Pgr*-Cre and *Ltf*-Cre models is the difference in the timing of Cre activation. *Pgr*-Cre becomes active approximately 10 days after birth ^30^, whereas *Ltf*-Cre is activated by estrogen at puberty ^31^. Thus, it is possible that defects in postnatal uterine adenogenesis contribute to the phenotypes observed in adult *Erbb3^d/d^* females. In our *Erbb3* mouse colony, 6 weeks of age corresponds to early puberty before the activation of *Ltf*-Cre. At this developmental stage, *Erbb3* transcripts were readily detected in both isolated epithelial and stromal cells from *Erbb3^f/f^* uteri (Supplementary Fig. 5). The majority of uterine glands in both *Erbb3^f/f^*and *Erbb3^d/d^* uteri have little or no branching at this stage (Fig. 3a and Supplementary Fig. 6). Quantitative analysis revealed comparable numbers of glands across branching categories between the two genotypes (Fig. 3b). Importantly, glandular FOXA2 expression was unaltered in *Erbb3^d/d^* uteri at this stage (Fig. 3c). Together, these results indicate that *Erbb3^d/d^* uteri display normal uterine gland morphology and FOXA2 expression prior to puberty, suggesting that the gland branching defects observed in adult *Erbb3^d/d^* females arise during or after puberty rather than from impaired postnatal adenogenesis. Because *Erbb3* deletion driven by *Pgr*-Cre has little or no impact on prepubertal uterine gland development, and the principal distinction between the *Pgr*-Cre and *Ltf*-Cre models lies in stromal *Erbb3* deletion after puberty, we infer that stromal *Erbb3* plays a critical role in regulating uterine gland branching and function.

**Fig. 3.**
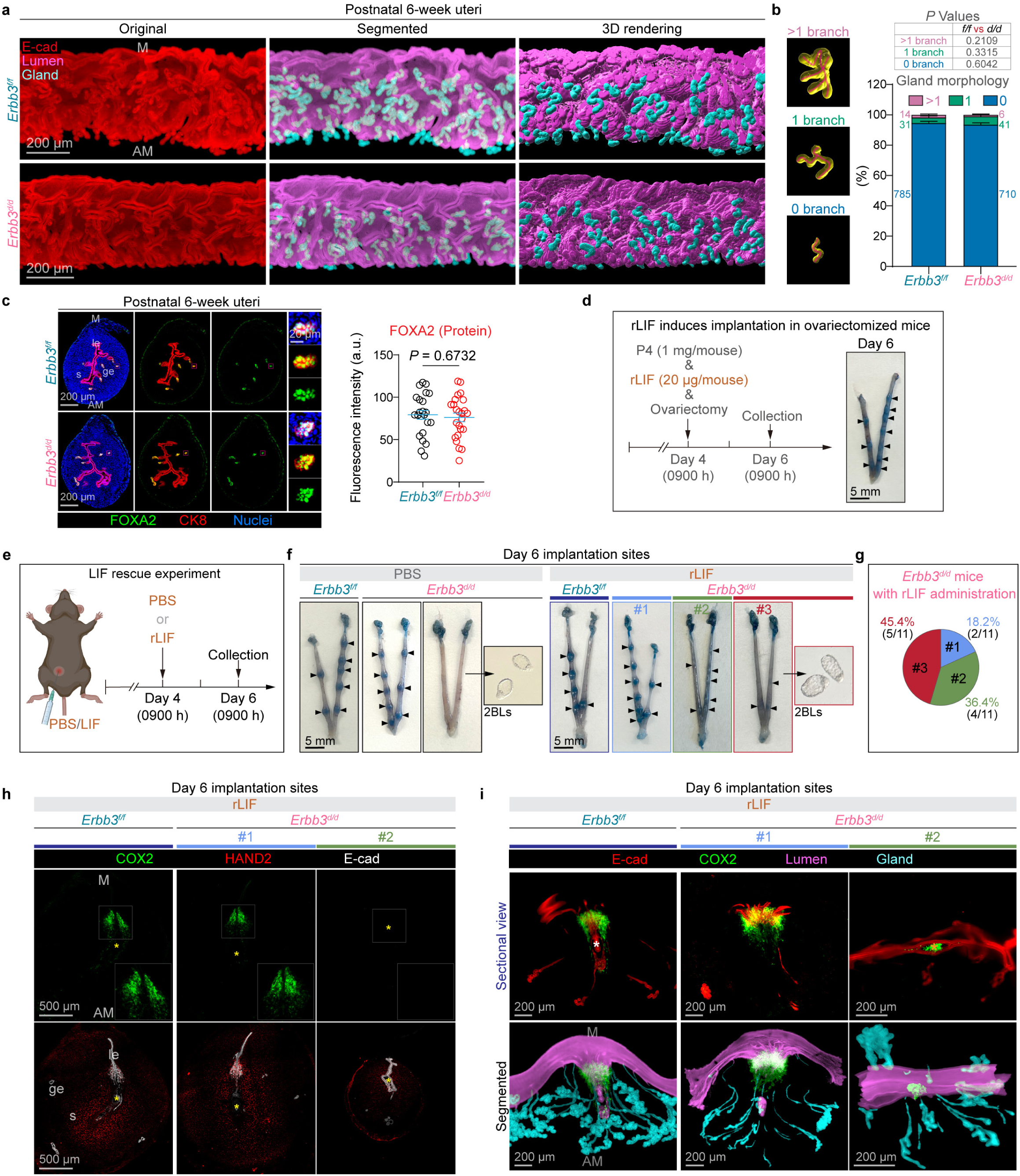
LIF administration failed to rescue implantation failure in *Erbb3^d/d^* females. **a** 3D imaging of postnatal 6-week uteri in *Erbb3^f/f^*(n = 6) and *Erbb3^d/d^* (n = 6) females. Original (E-Cadherin), segmented, and 3D rendering of 6-week uteri. Scale bars, 200 μm. **b** Quantification of gland branches in 6-week *Erbb3^f/f^*(n = 6) and *Erbb3^d/d^* (n = 6) females. Data are represented as mean ± SEM. *P* values were determined by a two-tailed Student’s *t* test. Source data are provided as a Source Data file. **c** Representative immunofluorescence images and quantification of FOXA2 expression in postnatal 6-week *Erbb3^f/f^*and *Erbb3^d/d^* uteri. The right panel shows the relative fluorescence intensity of FOXA2 within the glandular epithelium. Scale bars: 200 μm (main images) and 20 μm (magnified views). Data are represented as mean ± SEM (n = 6 mice per genotype). Statistical significance was assessed using a two-tailed Student’s *t* test. Source data are provided as a Source Data file. **d** LIF induces implantation in ovariectomized mice. A schematic illustrating the experimental procedure. Ovariectomized mice were subcutaneously injected with progesterone (P_4_, 1 mg/mouse) and intraperitoneally injected with recombinant LIF protein (rLIF, 20 μg/mouse) at 0900 h on Day 4. Uteri were collected at 0900 h on day 6. Arrowheads indicate implantation sites, confirming that rLIF can induce implantation. **e** A graphical diagram of the LIF rescue experiment. Created in BioRender. Li, B. (2026) https://BioRender.com/ucyr4o5. Mice were intraperitoneally injected with PBS or recombinant LIF (rLIF, 20 μg/mouse) at 0900 h on day 4, and uteri were collected at 0900h on day 6. **f** Representative images showing implantation sites on day 6 in *Erbb3^f/f^* and *Erbb3^d/d^*uteri after PBS or LIF treatment. Arrowheads indicate implantation sites. A boxed inset shows blastocysts recovered from the flushing from *Erbb3^d/d^* uteri. **g** Quantification of implantation outcomes in LIF-treated *Erbb3^d/d^* females (n = 11). **h** Immunofluorescence staining of day 6 implantation sites for COX2, HAND2, and E-cadherin (E-cad) of *Erbb3^f/f^* and *Erbb3^d/d^* uteri. Asterisks indicate the location of embryos. le, luminal epithelium; ge, glandular epithelium; s, stroma; M, mesometrial pole; AM, antimesometrial pole. Scale bars: 500 μm. **i** Sectional views (top) and 3D imaging (bottom) of day 6 implantation sites from LIF-treated *Erbb3^f/f^* and *Erbb3^d/d^* females. Original (E-cadherin and COX2), segmented, and 3D rendering of day 6 implantation sites in *Erbb3^f/f^*and *Erbb3^d/d^* females. Asterisks indicate the location of blastocysts. M, mesometrial pole; AM, antimesometrial pole.

### LIF administration failed to rescue implantation failure in *Erbb3^d/d^* females

Since *Lif* expression is reduced in *Erbb3^d/d^* glands on day 4, we examined whether exogenous LIF could rescue embryo implantation. After confirming the bioactivity of recombinant LIF protein (Fig. 3d) ^12^, we administered LIF or a PBS vehicle control intraperitoneally to *Erbb3^f/f^* and *Erbb3^d/d^* females on day 4 and assessed implantation on day 6 (Fig. 3e). As expected, PBS-treated *Erbb3^f/f^*females exhibited normal implantation, whereas a subset of PBS-treated *Erbb3^d/d^* females showed implantation failure, confirming that the vehicle alone does not alter the genotype-specific phenotypes (Fig. 3f). Administration of LIF had no detectable effect on implantation outcomes in *Erbb3^f/f^* females. In contrast, in the LIF-treated *Erbb3^d/d^* females (n = 11), only a small fraction (2 of 11, ∼18.2%) exhibited properly formed implantation sites (group #1). Around 1/3 of them (4 of 11) displayed overtly small, compromised implantation sites with faint blue bands (group #2). The remaining 5 females (5 of 11, ∼45.4%) had implantation failure with little or no detectable blue bands (group #3) (Fig. 3e-g).

To further investigate the quality of implantation in these females, we examined the expression of COX2 and HAND2, two established markers of implantation progression and decidualization ^32^. COX2 is initially expressed on the anti-mesometrial side of the implantation chamber on day 5 of pregnancy and subsequently shifts toward the mesometrial side as decidualization proceeds ^33^. COX2 signals were detected in implantation sites from *Erbb3^f/f^* controls and group #1 *Erbb3^d/d^* implantation sites (Fig. 3h). In contrast, little COX2 signals were detected in group #2 *Erbb3^d/d^* implantation sites. Additionally, the domain of HAND2 expression was reduced in group #2 implantation sites as compared to those of control and group #1 (Fig. 3h). COX2 expression patterns were also confirmed using 3D imaging. In group #2 *Erbb3^d/d^*implantation sites, embryos remained trapped within the luminal epithelium and displayed COX2 expression surrounding the embryo in a pattern resembling a typical day 5 implantation site (Fig. 3i), indicating a failure to progress beyond the initial attachment stage. Nonetheless, uterine glands in *Erbb3^d/d^* females from both group #1 and group #2 continued to exhibit reduced branching. Together, these observations indicate that restoration of LIF alone is insufficient to rescue implantation progression and decidualization in *Erbb3^d/d^* uteri.

### Uterine *Erbb3* deficiency impairs stromal growth factor networks required for uterine gland branching

Given the gland branching defects are observed in *Erbb3^d/d^*, but not *Erbb3^epi/epi^* uteri, we hypothesize that the stromal environment lacking ERBB3 signaling impairs normal gland branching. Our previous study demonstrated that uterine glands undergo progressive branching during the preimplantation period ^17^. By the evening of day 3 of pregnancy, a subset of glandular epithelial cells begins to express *Prss29* and becomes competent to respond to E_2_ stimulation for subsequent LIF production. Therefore, we selected day 3 of pregnancy to examine the stromal contribution to gland branching at a stage when glands are actively undergoing branching morphogenesis. To globally investigate the cellular composition and transcriptional landscape of the uteri, we performed single-cell RNA sequencing (scRNA-seq) from *Erbb3^f/f^* and *Erbb3^d/d^* uteri on day 3 using the 10x Genomics Chromium Flex (Flex Gene Expression) platform (Fig. 4a). *Erbb3^f/f^* and *Erbb3^d/d^* uteri have comparable cell populations (Fig. 4b), and an unsupervised cell clustering identified 11 distinct cell populations, including luminal epithelial (Le_1, Le_2, Le_3, and Le_4), glandular epithelial (Ge), stromal (Str), mesothelial (Meso), endothelial (Endo), perivascular (Peri), immune (Immune_1 and Immune_2) subsets (Fig. 4c). Each cluster was annotated based on canonical marker gene expression (Fig. 4d), and transcriptional similarities among clusters were assessed by correlation analysis (Fig. 4e). Cell-type proportion analysis also showed fewer glandular epithelial and stromal cells, with a relative increase in luminal epithelial cells in *Erbb3^d/d^* uteri compared to controls (Fig. 4f, g), suggesting a dysregulated stromal-epithelial balance in *Erbb3^d/d^*uteri.

**Fig. 4.**
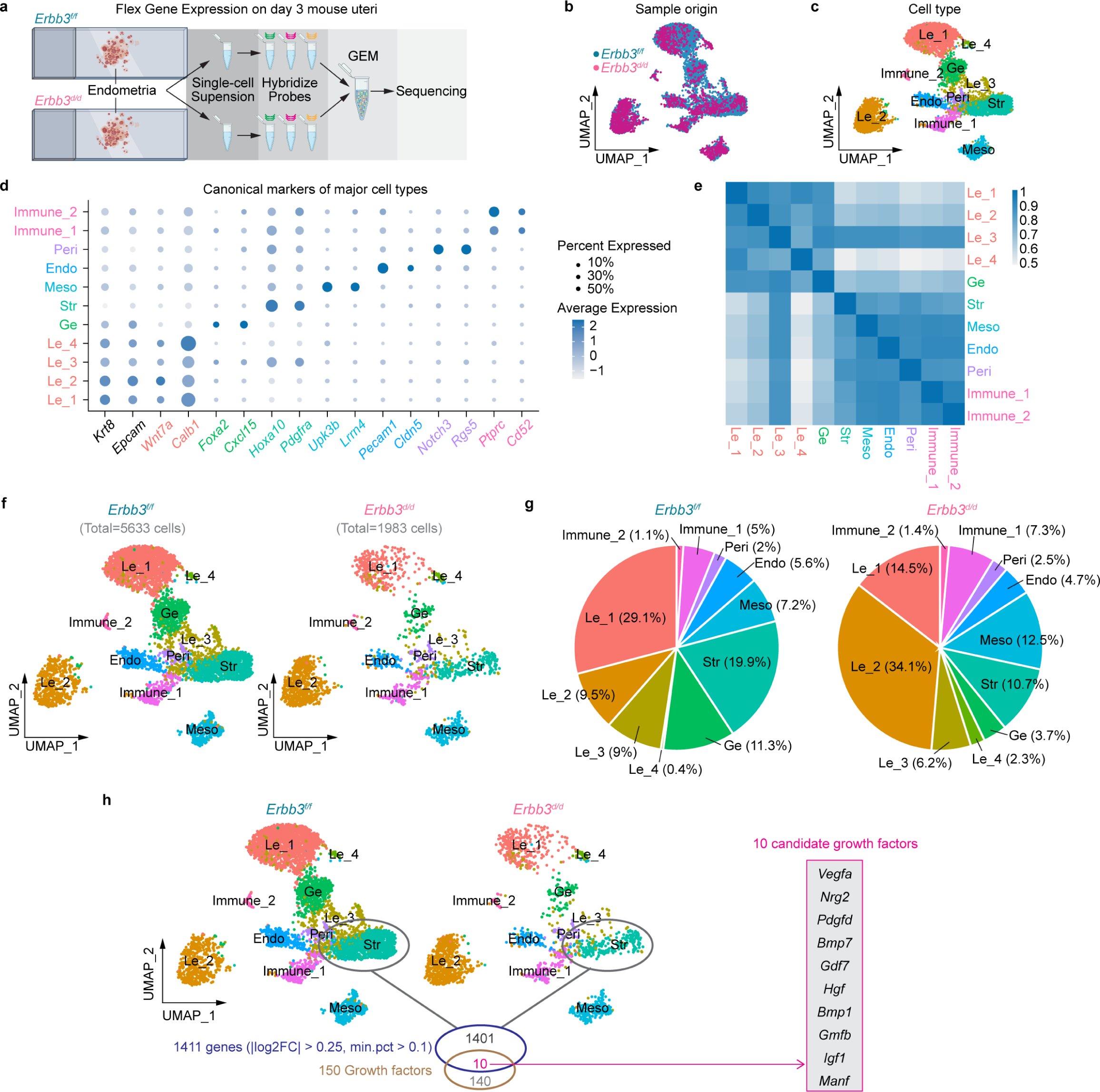
Single-cell transcriptomic profiling of day 3 *Erbb3^f/f^* and *Erbb3^d/d^* mouse uteri. **a** Schematic overview of the experimental workflow. Uteri from *Erbb3^f/f^* and *Erbb3^d/d^* females were collected on day 3 of pregnancy. Created in BioRender. Li, B. (2026) https://BioRender.com/danl3td. **b** UMAP plot showing distribution of cells by genotype: *Erbb3^f/f^* and *Erbb3^d/d^*. **c** UMAP plot showing clustering and annotation of major cell types. **d** Dot plot of canonical marker genes used for cell type identification. Dot size indicates the percentage of expressing cells, and color intensity represents average expression level. **e** Correlation heatmap showing transcriptional similarities among cell clusters. **f** UMAP plots of *Erbb3^f/f^* (left) and *Erbb3^d/d^* (right) cells, colored by cell type. **g** Pie charts showing the proportion of each cell type in *Erbb3^f/f^* (left, total 5633 cells) and *Erbb3^d/d^* (right, total 1983 cells) samples. **h** UMAP visualization of single-cell transcriptomes from day 3 *Erbb3^f/f^*and *Erbb3^d/d^* uteri, with the stromal cell (Str) population highlighted (circled). Differential expression analysis between *Erbb3^f/f^*and *Erbb3^d/d^* stromal cells (thresholds |log2(Fold Change)| > 0.25, min.pct > 0.1) identified 1,411 differentially expressed genes. Cross-referencing these with a curated growth factor gene set (GO:0008083; 150 genes) yielded 10 overlapping candidate growth factors, which are listed on the right.

Differential expression analysis of stromal cells from *Erbb3^f/f^*and *Erbb3^d/d^* uteri identified 1,411 genes meeting the specified thresholds (|log2FC| > 0.25, min.pct > 0.1). Cross-referencing these genes with a curated growth factor gene set (GO:0008083; 150 unique factors) revealed 10 candidate growth factors that were altered in *Erbb3^d/d^*stroma, including *Vegfa, Nrg2, Pdgfd, Bmp7, Gdf7, Hgf, Bmp1, Gmfb, Igf1,* and *Manf* (Fig. 4h). These scRNA-seq data indicate that uterine deletion of *Erbb3* disrupts stromal growth factor expression, potentially compromising stromal-epithelial communication during gland branching.

### Bulk transcriptomic analysis of isolated stroma confirms *Igf1* as a key ERBB3-dependent factor

To confirm the candidate growth factors identified by scRNA-seq analysis (Fig. 4h), uterine stromal cells were isolated from *Erbb3^f/f^* and *Erbb3^d/d^* females on day 3 of pregnancy and subjected to bulk RNA sequencing. Principal component analysis showed that most *Erbb3^d/d^* samples segregated from *Erbb3^f/f^*controls, although one *Erbb3^d/d^*sample clustered with the control group. This heterogeneity is consistent with our observation of incomplete phenotype penetrance in *Erbb3^d/d^*females (Supplementary Fig. 7a) and was further supported by sample-to-sample correlation analysis (Supplementary Fig. 7b). Quality control assessment of expression distributions demonstrated well-normalized raw count data across all eight samples, supporting the robustness of downstream analyses (Supplementary Fig. 7c).

Given the gland branching defects observed in *Erbb3^d/d^* uteri, we next examined whether established genes governing gland development were globally altered. Gene Set Enrichment Analysis (GSEA) revealed widespread changes in genes associated with “Gland Morphogenesis” following *Erbb3* deletion (Fig. 5a). Differential expression analysis identified 252 differentially expressed genes (DEGs), with 52 genes upregulated and 200 genes downregulated in *Erbb3^f/f^*stromal cells compared with *Erbb3^d/d^* stromal cells (Fig. 5b, c). Over-representation analysis of these DEGs showed strong enrichment for pathways related to reproductive system development, glandular and branching morphogenesis, and growth factor regulation (Fig. 5d), consistent with a requirement for stromal ERBB3 signaling in uterine gland development. We then examined individual genes within the enriched categories of “Morphogenesis of a branching epithelium” (Fig. 5e), “Gland morphogenesis” (Fig. 5f), and “Growth factors” (Fig. 5g). *Igf1* was identified as the top candidate gene among all three categories. Notably, among the 10 candidate growth factors identified by scRNA-seq (Fig. 4h), *Igf1* again emerged as the most significantly altered gene in the bulk RNA-seq dataset (Fig. 5h, i). To assess the spatial expression of *Igf1*, we performed fluorescence *in situ* hybridization on day 3 uterine sections, which revealed robust stromal *Igf1* expression in *Erbb3^f/f^* uteri that was significantly reduced in *Erbb3^d/d^* uteri (Fig. 5j). In addition, we confirmed reduced FOXA2 expression in *Erbb3^d/d^* uteri (Fig. 5j), consistent with impaired glandular development.

**Fig. 5.**
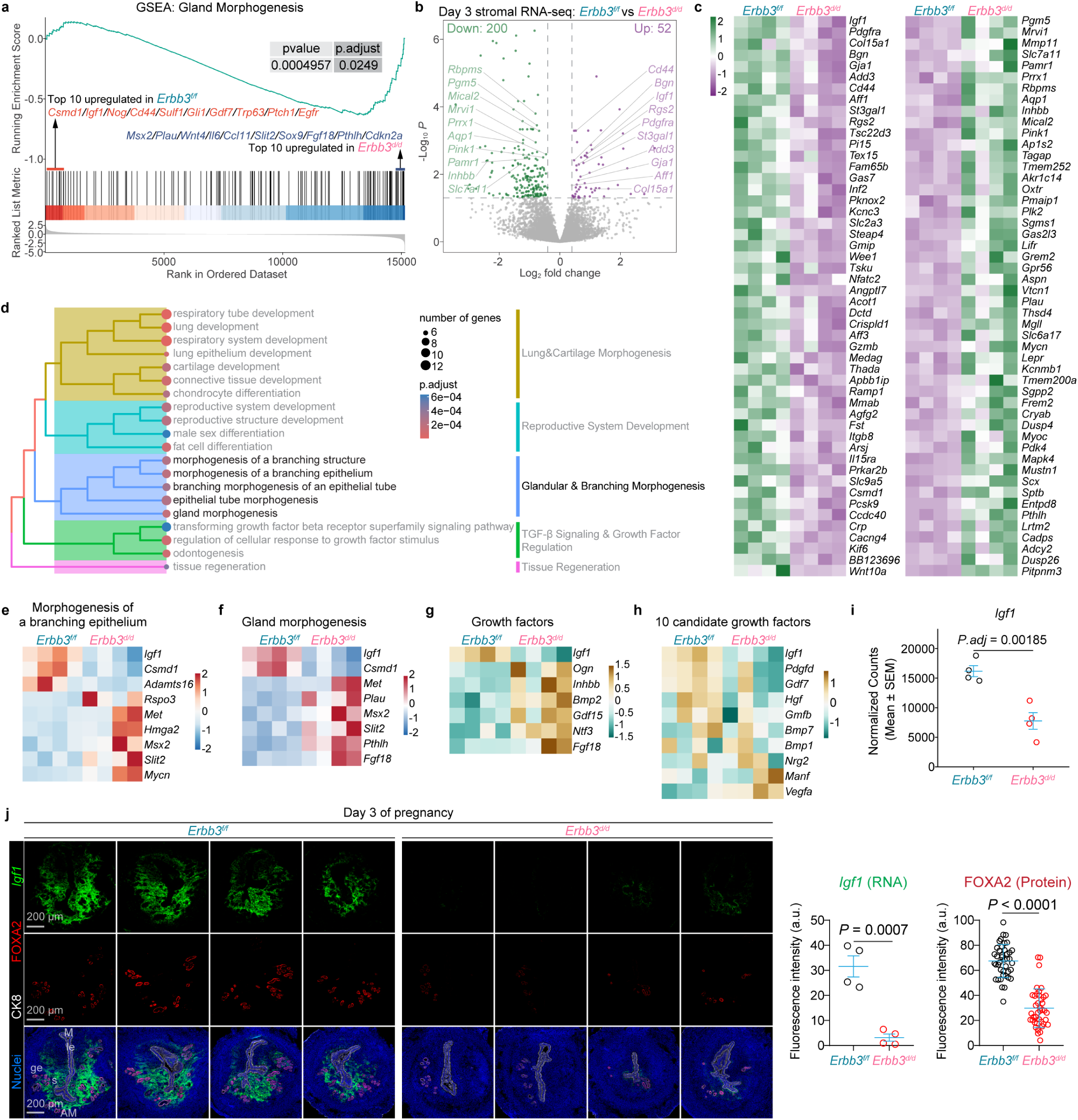
Uterine *Erbb3* deficiency impairs stromal growth factor networks. **a** GSEA Plot of gland morphogenesis (GO:0022612). Gene Set Enrichment Analysis (GSEA) was performed using the full transcriptomic profile of 15,151 detected genes, ranked by log2(Fold Change) from the most upregulated in the *Erbb3^f/f^* group to the most upregulated in the *Erbb3^d/d^* group. The enrichment score (green line) demonstrates a significant enrichment of the gland morphogenesis program (*p* = 0.0005, *p.adjust* = 0.0249, NES = −1.66). **b** Volcano Plot of Differentially Expressed Genes (DEGs). Statistical significance is plotted against fold change for all detected genes. Significant DEGs were defined by an adjusted p-value (*p.adjust*) < 0.05 and a |log2(Fold Change)| > 0.4. Purple and green points indicate genes significantly higher in the *Erbb3^f/f^* and *Erbb3^d/d^* groups, respectively (*p.adjust* < 0.05, |log2(Fold Change)| > 0.4); representative top 10 genes from each group are labeled. **c** Heatmap of the top 50 DEGs enriched in *Erbb3^f/f^* uteri and the top 50 DEGs enriched in *Erbb3^d/d^* uteri (excluding predicted genes with ‘Rik’ suffix), displaying normalized expression (Z-score). Genes are ranked by their expression levels within the respective group. Rows represent individual genes; columns represent biological replicates (n = 4 per group). **d** Treeplot of Functional Clustering. The treeplot illustrates the functional grouping of the top 20 most significant Biological Processes derived from Over-Representation Analysis (ORA). Input genes for this analysis were DEGs filtered by adjusted p-value (*p.adjust*) < 0.05 and |log2(Fold Change)| > 1. Enriched terms were clustered into five major functional modules based on pairwise semantic similarity. Each node represents a GO term, where the size reflects the gene count and the color indicates the statistical significance (adjusted p-value). **e** Heatmap showing normalized expression (*Z*-score) of DEGs (*p.adjust* < 0.05) enriched for “Morphogenesis of a Branching Epithelium” (GO:0061138). Genes are ranked by baseline expression in the *Erbb3^f/f^* group. **f** Heatmap of representative DEGs enriched for “Gland Morphogenesis” (GO:0022612). Color scale represents row-scaled *Z*-score (n = 4 biological replicates/group). **g** Heatmap of significant growth factors (GO:0008083; *p.adjust* < 0.05), sorted to display transcriptomic differences between *Erbb3^f/f^* and *Erbb3^d/d^*groups. **h** Heatmap of 10 candidate growth factors. Normalized expression (Z-score) of 10 manually curated growth factor candidates (*Vegfa*, *Nrg2*, *Pdgfd*, *Bmp7*, *Gdf7*, *Hgf*, *Bmp1*, *Gmfb*, *Igf1*, *Manf*) is displayed across all biological replicates (n = 4 per group). Genes are ranked by log2(Fold Change) from the most upregulated in the *Erbb3^f/f^* group (top) to the most upregulated in the *Erbb3^d/d^* group (bottom). Color scale represents row-scaled Z-score. **i** Normalized bulk RNA-seq counts of *Igf1* in *Erbb3^f/f^* and *Erbb3^d/d^* uteri on day 3 of pregnancy. Data are represented as mean ± SEM. P.adj value was determined by DESeq2 Wald test with Benjamini-Hochberg correction. Source data are provided as a Source Data file. **j** Validation of *Igf1* and FOXA2 expression in day 3 *Erbb3^f/f^* (n = 4) and *Erbb3^d/d^*(n = 4) uteri. Left: Co-staining of *Igf1*, FOXA2, and CK8. Scale bars, 200 μm. Right: Relative fluorescence intensity quantification of *Igf1* and FOXA2. Fluorescence intensity (a.u.) was quantified following the method described in the Methods section. le, luminal epithelium; ge, glandular epithelium; s, stroma; M, mesometrial pole; AM, antimesometrial pole. Data are presented as mean ± SEM. *P* values were determined by a two-tailed Student’s *t*-test. Source data are provided as a Source Data file.

### Stromal IGF1-epithelial IGF1R paracrine signaling as a candidate mediator of ERBB3 function

To independently confirm the findings from our scRNA-seq and bulk RNA-seq analyses, we reanalyzed a publicly available scRNA-seq dataset of day 3 pregnant mouse uteri (GSA: CRA011266) ^34^. Cells were reclustered by consolidating previously defined subpopulations into eight major cell types, including glandular epithelium (GE), luminal epithelium (LE), stromal cells (Str), and immune cells (Immune) (Supplementary Fig. 8a-c). We then assessed intercellular communication using ligand-receptor interaction analysis with CellChat. Collagen-mediated extracellular matrix signaling dominated the global communication network, with stromal cells serving as the primary signaling source (outgoing strength: 10.529), while LE and GE cells were the major signal recipients (incoming strengths: 7.503 and 6.307, respectively) (Fig. 6a-c).

**Fig. 6.**
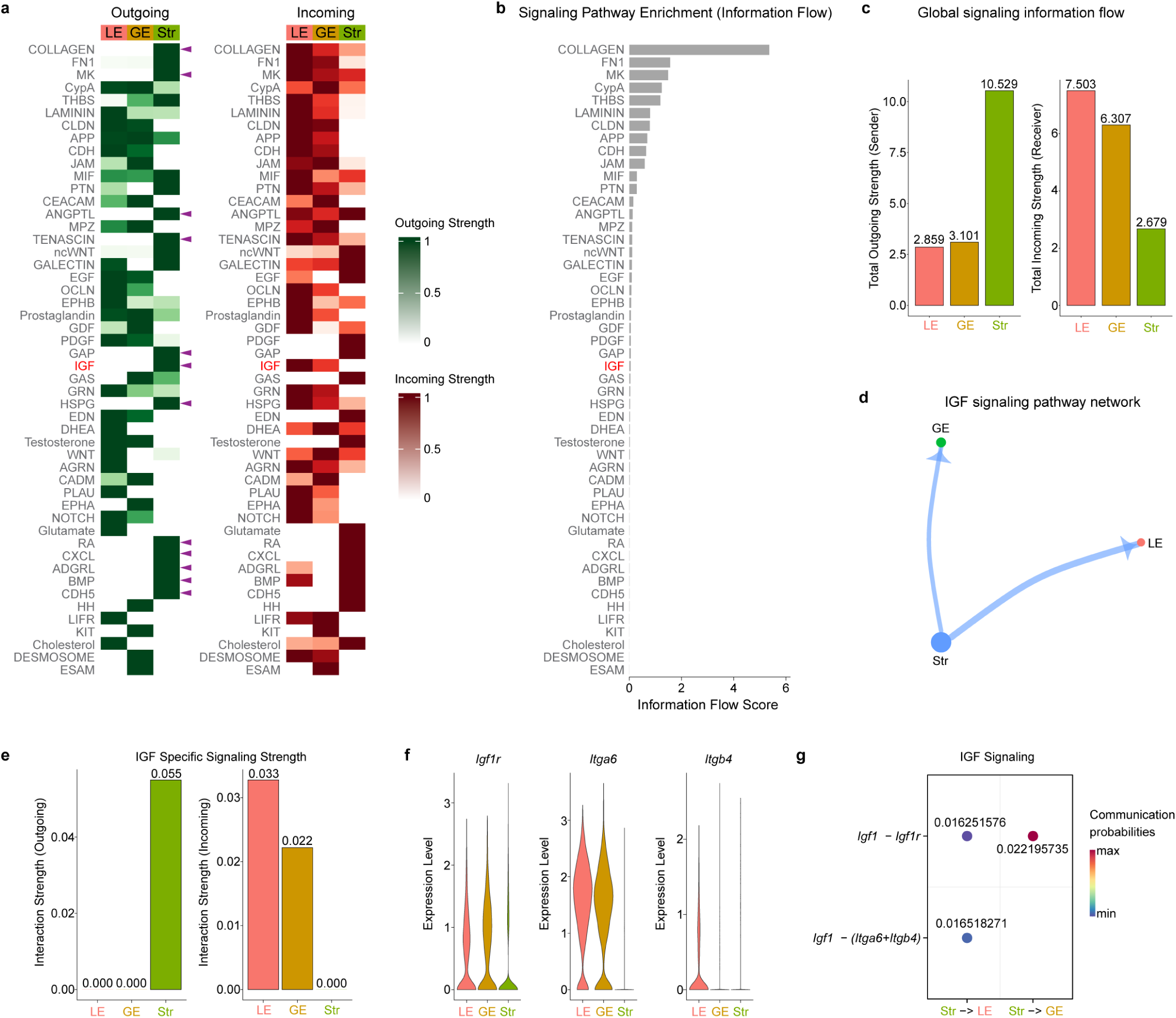
Single-cell transcriptomic analysis identifies a unidirectional stromal-to-epithelial *Igf1-Igf1r* signaling axis. **a** Heatmaps displaying global outgoing (sender) and incoming (receiver) signaling patterns across the LE, GE, and Str compartments. Purple arrowheads denote the 12 signaling networks exclusively originating from the stromal compartment. **b** Bar graph ranking the relative information flow (signal network probability) of detected pathways within the uterine microenvironment, highlighting the IGF pathway. **c** Quantification of total global communication strength, establishing the stroma as the dominant signal sender and the epithelia (LE and GE) as the primary receivers. Source data are provided as a Source Data file. **d** Hierarchical network topology of the IGF signaling pathway, illustrating a strictly unidirectional signal flow from the stroma to the LE and GE. **e** IGF-specific interaction strength analysis, confirming exclusive outgoing signals originating from the stroma and incoming signals strictly received by the epithelia. Source data are provided as a Source Data file. **f** Violin plots demonstrating the enriched expression of the primary IGF receptor (*Igf1r*) and its co-receptors (*Itga6*, *Itgb4*) in the epithelial compartments compared to the stroma. **g** Bubble plot detailing ligand-receptor interaction probabilities, confirming significant *Igf1-Igf1r* and *Igf1-(Itga6+Itgb4)* communication directed from the stroma to both LE and GE. Source data are provided as a Source Data file.

IGF signaling was again identified as one of the top stromal-to-epithelial communication pathways and exhibited a strictly unidirectional paracrine pattern (Fig. 6b, d). Specifically, stromal cells functioned as the exclusive source of IGF signals, transmitting exclusively to epithelial compartments (Fig. 6e). Gene expression mapping further supported this directionality, showing that the primary receptor *Igf1r* and its co-receptors *Itga6* and *Itgb4* ^35^ were highly enriched in epithelial populations on day 3 of pregnancy (Fig. 6f). Consistent with these findings, ligand receptor interaction analysis revealed significant communication probabilities for the *Igf1-Igf1r* axis directed from stromal cells to both luminal epithelium (0.016) and glandular epithelium (0.022) (Fig. 6g). Together, these analyses identified *Igf1-Igf1r* paracrine signaling as a strong candidate pathway through which stromal ERBB3 signaling influences uterine gland differentiation.

### Uterine deletion of *Igf1r* phenocopies the gland branching defects of *Erbb3*-deficient uteri

To validate the functional relevance of the IGF1-IGF1R paracrine pathway in gland branching, we examined uterine morphology in *Igf1r^f/f^ Pgr^Cre/+^* females on day 4 of pregnancy. Three-dimensional imaging revealed that uterine-specific deletion of *Igf1r* resulted in a significant reduction in gland branching compared with *Igf1r^f/f^ Pgr^+/+^* control mice (Fig. 7a, b and Supplementary Fig. 9). Quantitative analysis showed that whereas most glands in control uteri displayed complex higher-order branching (>1 branch), over half of the glands in *Igf1r^f/f^ Pgr^Cre/+^* uteri lacked branches (0 branch), remaining as primitive, elongated stems (Fig. 7c). These findings are consistent with previous reports demonstrating a significant reduction in the number of implantation sites on day 5 of pregnancy in *Igf1r^f/f^ Pgr^Cre/+^* females ^36^. Despite this pronounced branching defect, *Igf1r* deletion did not affect expression of the glandular marker FOXA2 (Fig. 7d), indicating that the IGF1-IGF1R paracrine pathway regulates gland branching through a FOXA2-independent mechanism.

**Fig. 7.**
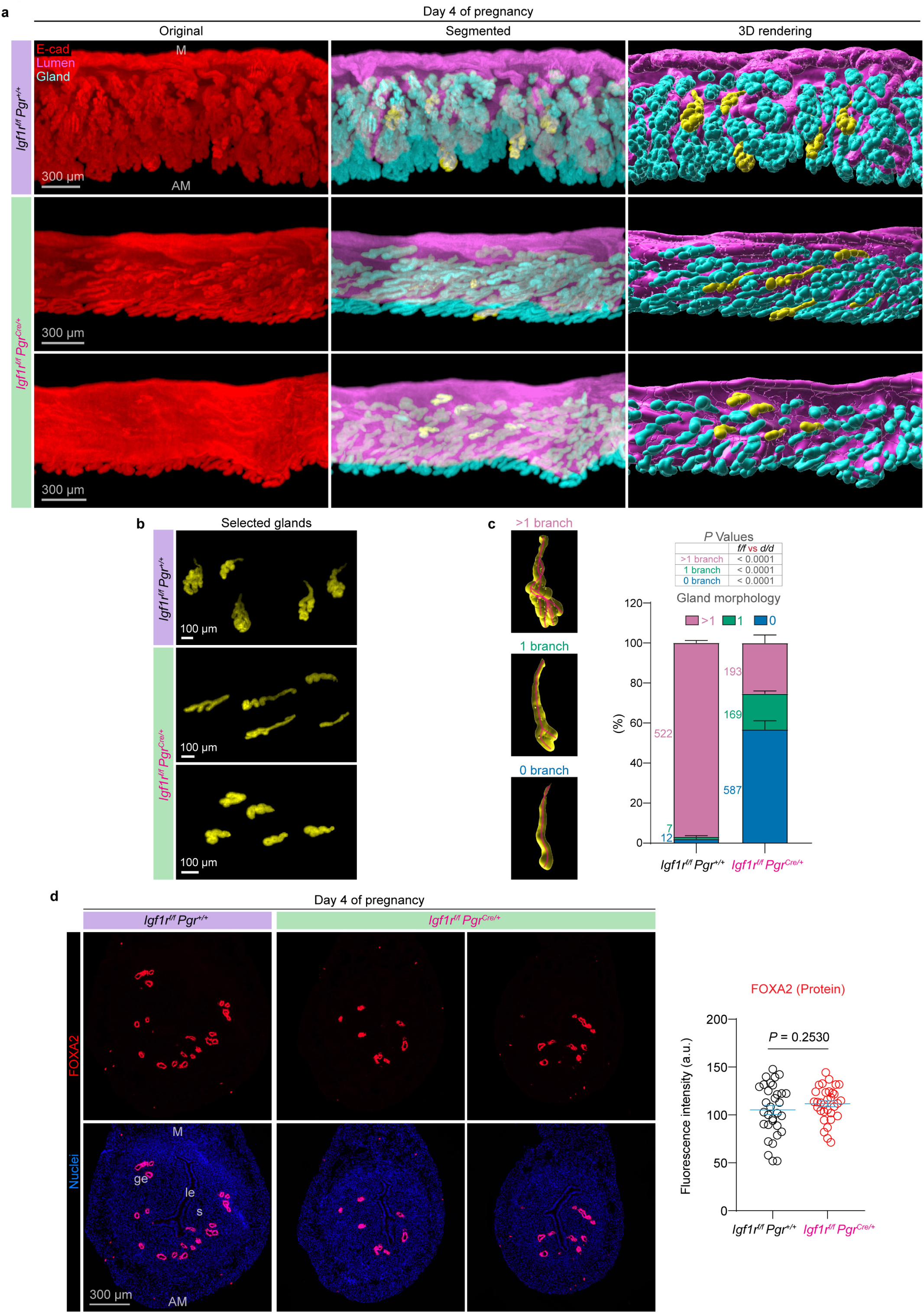
Uterine deletion of *Igf1r* impairs gland branching morphogenesis. **a** 3D whole-mount imaging of day 4 pregnant uteri from *Igf1r^f/f^ Pgr^+/+^* and *Igf1r^f/f^ Pgr^Cre/+^* females. Original E-cadherin, segmented, and 3D rendering. M, mesometrial pole; AM, antimesometrial pole. Scale bars, 300 μm. **b** High-magnification 3D reconstructions of selected glands extracted from **a**. Scale bars, 100 μm. **c** Quantification of gland branches from *Igf1r^f/f^ Pgr^+/+^* (n = 4), *Igf1r^f/f^ Pgr^Cre/+^* (n = 6) females on day 4 of pregnancy. Data are represented as mean ± SEM. *P* values were determined by a two-tailed Student’s *t* test. Source data are provided as a Source Data file. **d** Representative immunofluorescence images and quantification of fluorescence intensity (a.u.) for FOXA2 of day 4 uteri from pregnant *Igf1r^f/f^ Pgr^+/+^* (n = 3) and *Igf1r^f/f^ Pgr^Cre/+^* (n = 3) mice. Scale bar, 300 µm. le, luminal epithelium; ge, glandular epithelium; s, stroma; M, mesometrial pole; AM, antimesometrial pole. Data are presented as mean ± SEM. *P* values were determined by a two-tailed Student’s *t*-test. Source data are provided as a Source Data file.

We next asked whether IGF1R activation is diminished in *Erbb3^d/d^* uteri, which exhibit reduced *Igf1* expression. Previous studies have shown that IGF1 induces phosphorylation of IGF1R, leading to activation of AKT via phosphorylation at Ser473 as a canonical downstream signaling event ^37^. Western blot analysis of whole uterine tissue collected on day 4 of pregnancy revealed reduced levels of p-AKT in *Erbb3^d/d^* uteri compared with *Erbb3^f/f^* controls (Supplementary Figs. 10a, b). To assess the cellular specificity of this defect, we performed immunostaining and found that p-AKT was broadly distributed across both stromal and epithelial compartments. Notably, p-AKT signals were markedly reduced in luminal and glandular epithelial cells of *Erbb3^d/d^* uteri on days 3 and 4 of pregnancy (Supplementary Figs. 10c, d). Together, these findings indicate that uterine deletion of *Erbb3* impairs IGF1R activation, resulting in reduced epithelial AKT signaling, and support a model in which stromal ERBB3 maintains *Igf1* production to drive epithelial IGF1R-AKT signaling essential for preimplantation gland branching.

## Discussion

Our current study demonstrates that EGF signaling is critical for uterine gland morphogenesis and differentiation. Uterine glands originate from the luminal epithelium ^38^, invaginate into the underlying stroma, and subsequently undergo elongation and branching. These morphogenetic processes are essential for establishing a functional glandular network critical for embryo implantation during mouse pregnancy ^38^. Multiple signaling pathways have been shown to play critical roles in early uterine adenogenesis, among which Wnt signaling is well characterized. Loss-of-function of *Wnt5a*, *Wnt7a*, or *Ctnnb* results in a glandless uterine phenotype ^39, 40, 41, 42^. Similarly, neonatal deletion of *Wnt4* in the uterus leads to a reduction in gland number ^43^. Neonatal uterine deletion of *Foxa2* or an adherens junction protein *Cdh1* also abolish adenogenesis ^5, 44^. However, detailed studies of uterine gland morphology were limited until the development of three-dimensional visualization methods for the uterus ^15, 16^. Two recent studies showed that post-pubertal deletion of *Foxa2* and prenatal epithelial deletion of *Esr1* result in branchless glands and compromised gland function ^17, 18^, suggesting that gland branching is essential for proper glandular function. Beyond these studies, our understanding of the signaling pathways that regulate uterine gland branching remains limited. Our current study identifies ERBB3-mediated EGF signaling as a key regulator of uterine gland branching and function, which advances our understanding of the molecular mechanisms underlying uterine gland branching.

An important caveat in comparing the *Pgr-*Cre and *Ltf-*Cre models is the difference in the timing of Cre activation. *Pgr-*Cre becomes active approximately 10 days after birth ^30^, whereas *Ltf-*Cre is activated by estrogen at puberty ^31^. Therefore, it is possible that defects in neonatal uterine adenogenesis in *Erbb3^d/d^* mice substantially contribute to the phenotypes observed in adulthood. To address this possibility, we compared uterine gland morphology in *Erbb3^d/d^* and *Erbb3^f/f^*mice at 6 weeks of age, immediately prior to the initiation of *Ltf*-Cre activity. In our *Erbb3* mouse colony, 6 weeks of age corresponds to early puberty, a developmental stage during which females have undergone minimal to no estrous cycles. As shown in Figs 3a and 3b, quantitative 3D imaging analysis revealed no significant differences in gland branch number between *Erbb3^d/d^*and *Erbb3^f/f^* uteri (P > 0.05 across all branching categories). In addition, expression of the glandular marker FOXA2 was comparable between genotypes (Fig. 3c). These findings suggest that ERBB3 signaling influences glandular FOXA2 expression only in the presence of ovarian hormones. Although the mechanism underlying the stage-specific regulation of FOXA2 by ERBB3 remains unclear, our data indicate that early postnatal uterine adenogenesis is not impaired in *Erbb3^d/d^* mice. Therefore, the gland branching defects observed after puberty are unlikely to arise from a pre-existing developmental abnormality and instead reflect a requirement for ERBB3 signaling during the post-pubertal period.

*Erbb3* expression in the stromal compartment plays a critical role in uterine gland branching. To date, the only mouse models reported to exhibit branchless uterine glands involve epithelial-specific deficiencies of *Foxa2* or *Esr1* ^17, 18^. These findings raise the question of whether uterine gland branching is regulated intrinsically by the glands themselves during the estrous cycle and early pregnancy. *Erbb3* is expressed in both uterine epithelial and stromal cells. *Erbb3^f/f^ Pgr^Cre/+^* females with deletion of *Erbb3* in both epithelia and stroma display compromised gland branching, whereas *Erbb3^f/f^ Ltf^Cre/+^*females with epithelial specific deletion of *Erbb3* show normal gland morphogenesis. Notably, pre-pubertal adenogenesis is not impaired in *Erbb3^f/f^ Pgr^Cre/+^* uteri. Collectively, these results suggest that loss of stromal *Erbb3* is required to drive the gland branching defects. However, we cannot conclude that stromal *Erbb3* deletion alone is sufficient to drive the defects, as direct evidence from a stroma-specific knockout model is currently lacking. *Amhr2*-Cre has been used to delete genes in the uterine stroma, but a recent study showed that *Amhr2*-Cre has Cre activity in early embryos resulting in global genetic modification ^45^; *Pdgfra*-Cre^ERT2^ has been used for uterine stromal lineage tracing experiments ^46^, but *Pdgfra*-Cre has activity in mesenchymal cells across multiple organs beginning at embryonic stages. Despite these limitations, our study highlights the essential role of stromal ERBB3 signaling in post-pubertal gland morphogenesis.

The stromal environment plays a critical role in shaping uterine glandular cell identity. *Foxa2* is expressed in uterine glands beginning at neonatal adenogenesis, and its gland-specific expression is maintained throughout adulthood ^5, 13, 14^. FOXA2’s faithful presence in uterine glands makes it a putative marker of uterine glands. In humans, dysregulation of endometrial FOXA2 is associated with several uterine pathologies, including complex atypical endometrial hyperplasia, uterine carcinoma, and endometriosis ^11, 47^. In mice, uterine deletion of *Foxa2* disrupts glandular function and compromises female fertility ^5, 13, 14^. In addition, we recently demonstrated that uterine glands progressively acquire branches from day 1 to day 4 of pregnancy, and uterine epithelial deletion of *Foxa2* (*Foxa2^f/f^ Ltf^Cre/+^*) impairs glandular branching ^17^. These findings indicate that FOXA2 is essential for maintaining uterine gland identity and proper gland morphogenesis. However, because uterine glandular epithelium originates from FOXA2-negative luminal epithelium, it remains unclear whether FOXA2 expression in uterine glands is autonomously maintained by glandular epithelial cells or regulated by signals from the surrounding stroma. In this study, we show that FOXA2 levels are reduced in adult, but not prepubertal, *Erbb3^d/d^* glands (Figs. 2h and 3c), suggesting that uterine glandular cell identity is compromised in the absence of post-pubertal EGF signaling. Nevertheless, it is still unknown whether loss of FOXA2 causes glandular epithelial cells to revert to a luminal epithelial fate, or whether these FOXA2-negative glandular cells persist in an intermediate or transitional state between glandular and luminal epithelial identities.

Exogenous LIF administration is insufficient to functionally rescue implantation failure in *Erbb3^d/d^* mice. In *Erbb3^d/d^*uteri, defects in gland branching and reduced FOXA2 expression in uterine glands ultimately result in impaired glandular function and diminished LIF production (Fig. 2). However, supplementation with exogenous LIF failed to rescue the implantation failure observed in *Erbb3^d/d^*females. These findings indicate that uterine *Erbb3* deficiency leads to multiple defects in uterine receptivity, including glandular LIF insufficiency. Consistent with this interpretation, the reduced stromal IGF1 levels in *Erbb3^d/d^* uteri further indicate that stromal cell function is also compromised. Because stromal, luminal epithelial, and glandular epithelial cells all play key roles in uterine receptivity, LIF supplementation alone is insufficient to overcome the implantation failure in *Erbb3^d/d^*females.

Stromal IGF1 mediates the impact of ERBB3 signaling on uterine gland morphogenesis. Our finding that stromal ERBB3 is essential for gland branching prompted the hypothesis that stromal ERBB3 influences glandular morphology through one or more secreted mediators. *Igf1* was identified as a strong candidate mediator by our scRNA-seq and bulk RNA-seq analyses intersecting with a secreted protein library. *In vivo*, endometrial cells can be influenced by two sources of IGF1: systemic circulating IGF1 and IGF made locally within the uterus. Circulating IGF1 is secreted predominantly by the liver. A previous study demonstrated that *Igf1^f/f^ Pgr^Cre/+^* females show decreased fertility ^48^, indicating that systemic IGF1 is unable to compensate for reduced local IGF1 levels in the uterus during pregnancy. *Igf1^f/f^ Pgr^Cre/+^* females have oviductal embryo retention and compromised preimplantation embryo development ^49^, which precludes their use for implantation studies. Gland branching and differentiation occur over several days, making functional rescue experiments particularly challenging. Moreover, IGF1 is a potent growth factor and systemic hormone *in vivo* (reviewed in ^50^). Repeated administration of exogenous IGF1 can induce marked physiological effects, including acute hypoglycemia, renal fluid retention, and increased capillary permeability. Given these broad mitogenic and metabolic actions, systemic IGF1 treatment would confound attempts to directly assess IGF1-mediated rescue of uterine gland defects.

Stromal IGF1 mediates ERBB3’s impact on gland morphogenesis via targeting epithelial IGF1R. Our cell-cell communication analysis using scRNA-seq data suggested that stromal IGF1 likely acts through its receptor IGF1R on epithelial cells. A recent study demonstrated that IGF1R is predominantly expressed in the uterine epithelium throughout early pregnancy ^36^, while *Igf1* ligand expression is mainly expressed in the stroma. The spatial proximity strongly supports that stromal IGF1 signals directly through epithelial IGF1R. In fact, uterine deletion of *Igf1r* leads to reduced gland branching similar to the phenotype of *Erbb3^d/d^* uteri. A comparable mechanism has been reported in the mammary gland, where IGF1 stimulation promotes epithelial expansion, while epithelial-specific *Igf1r* deficiency leads to branching defects ^51^. These findings collectively support a model in which IGF1-IGF1R signaling mediates stromal-epithelial communication during gland development. IGF1 can also target the insulin receptor (INSR), which is expressed in the preimplantation uterus. However, recent studies showed that *Insr^f/f^ Pgr^Cre/+^* females have numbers of implantation sites comparable to those of control females on day 6 of pregnancy, whereas *Igf1r^f/f^ Pgr^Cre/+^*females display a significant reduction in implantation sites ^36, 52^. These findings suggest that uterine IGF1R plays a more critical role than INSR during early pregnancy. Although we cannot exclude a contributory role for INSR in mediating IGF1 signaling, the glandular morphogenetic defects observed in *Igf1r^f/f^ Pgr^Cre/+^* uteri demonstrate that IGF1R is a key mediator of uterine gland branching.

Surprisingly, FOXA2 expression is comparable between *Igf1r^f/f^ Pgr^Cre^*^/+^ and control uterine glands, whereas it is markedly reduced in *Erbb3^d/d^* glands. This finding suggests that ERBB3 regulates glandular branching through both a FOXA2-dependent and a FOXA2-independent mechanism mediated by the IGF1-IGF1R pathway (Fig. 8). Nevertheless, several important questions remain unresolved. The mechanism by which ERBB3 regulates IGF1 expression is unclear. In addition, how ERBB3 controls glandular FOXA2 expression and how IGF1R activation promotes gland branching are not fully understood. Addressing these questions will require further investigation.

**Fig. 8.**
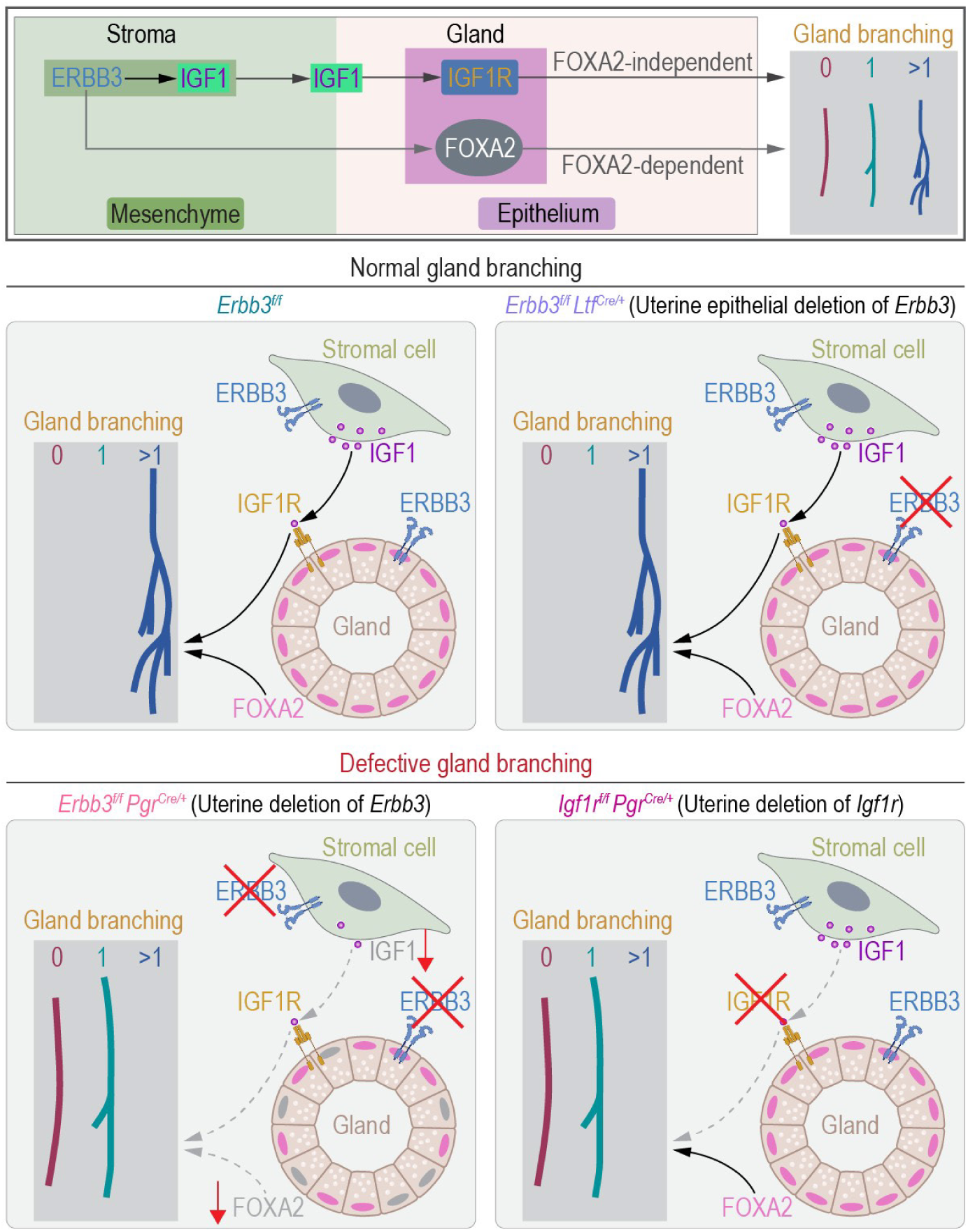
A schematic model illustrating how *Erbb3*-*Igf1*-*Igf1r* signaling regulates uterine gland branching. In control (*Erbb3^f/f^*) or epithelial-*Erbb3*-deleted (*Erbb3^f/f^ Ltf^Cre/+^*) uteri, stromal ERBB3 promotes complex gland branching via two parallel mechanisms: maintaining glandular FOXA2 expression and driving the paracrine IGF1-IGF1R signaling axis. Ablating whole-uterus *Erbb3* (*Erbb3^f/f^ Pgr^Cre/+^*) impairs both glandular FOXA2 and stromal IGF1, arresting branching morphogenesis. Deletion of the IGF1 receptor (*Igf1r^f/f^ Pgr^Cre/+^*) blocks IGF1 signal reception without altering FOXA2 levels, and disruption of this IGF1-IGF1R paracrine axis partially phenocopies the glandular branching failure. Created in BioRender. Li, B. (2026) https://BioRender.com/btxy3gu.

## Methods

### Mice

*Erbb3^f/f^* ^53^ mice were crossed with *Pgr^Cre/+^* ^30^ or *Ltf^Cre/+^* ^31^ mice to generate *Erbb3^f/f^ Pgr^Cre/+^* (*Erbb3^d/d^*) and *Erbb3^f/f^ Ltf^Cre/+^* (*Erbb3^epi/epi^*) mouse lines. *Erbb3^d/d^*, *Erbb3^epi/epi^* and their control littermates were housed in the animal care facility at Cincinnati Children’s Hospital Medical Center according to the National Institute of Health and institutional guidelines for laboratory animals. All protocols were approved by the Cincinnati Children’s Animal Care and Use Committee. Mice were provided with irradiated Laboratory Rodent Diet 5R53 and autoclaved water ad libitum. These mice were housed under a 12:12 hr light:dark cycle. *Pgr^Cre^*^/+^ mice were originally provided by Francesco DeMayo (National Institute of Environmental Health Science, Research Triangle Park, NC) and John B. Lydon (Baylor College of Medicine, Houston, TX). *Erbb3* floxed mice were provided by David Threadgill (Texas A&M University, College Station, TX). *Igf1r^f/f^ Pgr^Cre/+^* mouse line ^36^ was obtained from Wenbo Deng (Xiamen University). At least three mice from each genotype were used for each individual experiment.

### Analysis of pregnancy events

Adult females from each genotype were randomly chosen and housed with a fertile WT male of choice overnight; the morning of finding a vaginal plug was considered successful mating (day 1 of pregnancy). Pregnancy in plug-positive mice was confirmed on day 4 by flushing one uterine horn with saline to identify blastocysts. For pregnant mice on days 5 and 6, 100 μl of 1% Chicago blue in saline was injected intravenously for four minutes to visualize implantation sites as blue bands. In the absence of blue bands, uterine horns were flushed to assess the presence of embryos.

### Fluorescence *In Situ* Hybridization (FISH)

FISH experiments were performed as previously described ^12^. Briefly, frozen sections (12-µm) were processed on the same slide for each probe. Following fixation in 4% paraformaldehyde and acetylation, slides were hybridized at 55°C with digoxigenin-labeled *Lif*, *Igf1*, or *Msx1* probes. Anti-DIG-peroxidase was applied onto hybridized slides following washing and peroxide quenching. The color was developed by a fluorescein Tyramide Signal Amplification kit according to the manufacturer’s instructions (PerkinElmer). Counterstaining was performed using a CK8 antibody to label epithelial cells (TROMA-I, DSHB) and a FOXA2 antibody to label glandular cells (8186S, Cell Signaling Technology). Nuclei were counterstained with Hoechst 33342 (2 µg/ml, H1399, Thermo Scientific). Images were acquired using a confocal microscope (Nikon Eclipse TE2000).

### Immunofluorescence (IF)

For immunofluorescence, 12-µm frozen sections from each genotype were co-mounted on single slides to standardize staining conditions. Sections were incubated with primary antibodies against ESR1 (sc-542, Santa Cruz; 1:300), PR (8757, Cell Signaling Technology; 1:300), Ki67 (RM-9106-S, Invitrogen; 1:300), HAND2 (AF3876, R&D systems; 1:300), Vimentin (5741, Cell Signaling Technology; 1:300), CK8 (TROMA-1, Developmental Studies Hybridoma Bank; 1:300), E-cadherin (13-1900, 1:300, Invitrogen), FOXA2 (8186s, Cell Signaling Technology; 1:200), or Phospho-Akt (Ser473) (9277, Cell Signaling Technology; 1:300). Signals were visualized using Alexa Fluor 488-, 594-, or 647-conjugated secondary antibodies (Jackson ImmunoResearch; 1:200). Nuclei were counterstained with Hoechst. Confocal images were acquired using a Nikon Eclipse TE2000 microscope. The mesometrial pole is the dorsal side of the uterus where blood vessels enter and the placenta forms; the antimesometrial pole is the ventral side where a blastocyst initially attaches. Histological sections were oriented and labeled accordingly to ensure consistent regional comparisons.

### Quantification of Fluorescence Intensity

Fluorescence intensity was quantified using ImageJ (NIH). Briefly, fluorescence images were opened and split into individual channels using the Split Channels function under the Image Color menu. The channel corresponding to the target signal was selected for analysis. For each image, regions of interest (ROIs) were manually delineated using a freehand selection tool, and the mean gray value within each ROI was quantified using the *Measure* function. All images within each comparison group were acquired using identical microscope settings, and fluorescence intensities are reported in arbitrary units (a.u.). The detailed step-by-step methods are illustrated in Supplementary Fig. 11. For quantification of *Lif* RNA and FOXA2 protein signals, fluorescence intensity was measured from each individual gland.

### Quantification of serum E_2_ and P_4_ levels

Sera were collected on day 4 of pregnancy (1100 h), and hormone levels were measured by enzyme immunoassay kits (Estradiol ELISA Kit, 501890 & Progesterone ELISA Kit, 582601, Cayman) as previously described ^31^.

### Whole-mount immunostaining for 3D imaging

The whole-mount immunostaining with 3DISCO tissue clearing was performed as previously described ^16^. Briefly, mouse uteri were fixed in Dent’s fixative (methanol:DMSO, 4:1) at −20 °C overnight and then washed with 100% methanol three times for 1 hour each. The samples were bleached with 3% H_2_O_2_ in methanol at 4 °C overnight. After three washes in 1% PBST for 1 hour each, the samples were blocked in 5% BSA and then incubated with anti-E-cadherin (13-1900, 1:200, Invitrogen) and/or anti-COX2 (12282s, 1:100, Cell Signaling Technology) in the same buffer at 4 °C on a rotor for 7 days. After incubation, the samples were washed with 1% PBST six times for 1 hour each and incubated with Alexa-conjugated 488, 594 or 647 antibodies (1:200, Jackson Immuno Research) in a light-proof box for 4 days at 4 °C. After six washes in 1% PBST at room temperature, the samples were dehydrated in 100% methanol for 30 minutes and then cleared in benzyl alcohol/benzyl benzoate (BABB) solution for at least 1 hour in a light-proof box.

### 3D imaging and processing

3D pictures were acquired using a Nikon FN1 Upright Microscope. Samples were laid on slides, covered with BABB, and enclosed by cover slips for confocal imaging using a 10X objective with 8 µm Z-stack. Image files generated using Nikon NIS-Elements were imported into Imaris software (version 10.2.0; Bitplane) for visualization and three-dimensional (3D) reconstruction. Tissue 3D structures were generated using the *Surface* tool. Specific tissue regions were isolated by manual segmentation with the *Surface* tool following a published protocol (“Segmentation Using Imaris 10.2.0” ^17^), and the *Mask* option was applied for subsequent pseudocoloring. Three-dimensional images were generated using the *Snapshot* tool.

### Isolation of mouse uterine epithelial and stromal cells

Epithelial and stromal cells were isolated from mouse uteri by sequential enzymatic digestion as described previously ^54^. Uteri were slit open longitudinally and cut into small fragments. For day 3 and day 4 pregnant uteri, tissue fragments were incubated with pancreatin (25 mg/mL; Sigma) and dispase (6 mg/mL; Gibco) for 30 min at 4 °C, followed by 20 min at room temperature and an additional 10 min at 37 °C. For uteri from 6-week-old adult mice, the same enzyme concentrations were used, but incubation was performed at 37 °C for 10 min. In both cases, luminal epithelial sheets were removed by gentle washing with HBSS and collected separately. The remaining tissue was then digested with type IV collagenase (0.5 mg/mL; Worthington) for 30 min at 37 °C to release stromal cells. Cell suspensions were filtered through a 70-μm nylon mesh to remove tissue clumps. The separated epithelial and stromal fractions were collected for RNA extraction and qPCR analysis. The purity of each fraction was confirmed by immunofluorescence staining for Vimentin (stromal cells), FOXA2 (glandular epithelium), and E-cadherin (all epithelium).

### RNA extraction and quantitative PCR (qPCR)

Total RNA was extracted from uterine tissues using TRIzol Reagent (Invitrogen, Cat# 15596026) according to the manufacturer’s instructions. RNA concentration and purity were assessed by measuring absorbance at 260 and 280 nm; samples with A260/A280 ratios between 1.8 and 2.0 were used for downstream analysis. Five hundred nanograms of total RNA were reverse transcribed into cDNA using the PrimeScript RT Reagent Kit (TaKaRa Bio, Cat# RR037A) following the manufacturer’s protocol. Quantitative PCR was performed on a StepOnePlus Real-Time PCR System (Applied Biosystems) using Fast SYBR Green Master Mix (Thermo Fisher Scientific). Reactions were run in triplicate with 1 μM primers. Relative gene expression was calculated using the 2⁻^ΔΔCt^ method and normalized to *Gapdh*. Primer sequences are listed in Supplementary Table 2.

### Western blotting

Protein extraction and Western blotting were performed as previously described ^55^. The primary antibodies used in this study included rabbit anti-ERBB3 Antibody (1:1000; 12708s, Cell Signaling Technology), rabbit anti-FOXA2 antibody (1:1000; 8186s, Cell Signaling Technology), rabbit anti-AKT Antibody (1:1000; 9272s, Cell Signaling Technology), rabbit anti-p-AKT (Ser473) Antibody (1:1000; 4060s, Cell Signaling Technology), and rabbit anti-GAPDH antibody (1:1000; 2118s, Cell Signaling Technology). Bands were visualized using an ECL Prime Western blotting detection system (GE Healthcare). GAPDH served as loading controls. Uncropped blot images are provided in Supplementary Fig. 12.

### Bulk RNA-sequencing analysis

Total RNA was extracted from day 3 mouse uterine stromal cells and subjected to paired-end RNA sequencing. Reads were aligned to the mouse reference genome (mm10) using HISAT2 with default settings. Alignments were converted to BAM and coordinate-sorted with samtools. To restrict analyses to uniquely mapped primary alignments, reads with NH:i:1 and without secondary/supplementary flags were retained. Gene-level read counts were obtained from the sorted BAM files using featureCounts (paired-end mode, chimeric/discordant pairs excluded) against a gene annotation (GTF) matching mm10. Differential expression analysis was performed in DESeq2 on raw counts; p values were adjusted by the Benjamini-Hochberg method, and genes with adjusted p < 0.05 were considered differentially expressed. Variance-stabilized counts were used for PCA and sample-to-sample correlation. Functional enrichment of DEGs was assessed using clusterProfiler for Gene Ontology biological processes.

### GSEA

Gene Set Enrichment Analysis (GSEA) was performed using the gseGO function in the clusterProfiler R package with the org.Mm.eg.db annotation database. Rather than restricting the analysis to differentially expressed genes, we used the full transcriptomic profile of all detected genes (∼15,000). Genes were ranked by their log2 fold change values derived from the DESeq2 analysis (*Erbb3^f/f^* versus *Erbb3^d/d^*). Enrichment was assessed against Gene Ontology Biological Process (GO:BP) gene sets, with gene set sizes constrained between 10 and 500 genes. P values were adjusted using the Benjamini-Hochberg method, and terms with adjusted P < 0.05 were considered significantly enriched. A permutation seed of 123 was used to ensure reproducibility. Core enrichment (leading-edge) genes were extracted for downstream pathway analysis.

### GEM-X Flex single-cell gene expression assay

Sample preparation was performed using the GEM-X Flex Sample Preparation v2 Kit (1000781, 10x Genomics) according to the manufacturer’s protocol. Endometrial tissues (∼25 mg) from *Erbb3^f/f^* and *Erbb3^d/d^* females were isolated by scraping the luminal surface of the uterus with a scalpel on a pre-chilled glass slide maintained on ice, thereby excluding the myometrial layer, finely minced, and fixed in Fixation Buffer B (nuclease-free water, Conc. Fix & Perm Buffer B, and 37% formaldehyde) for 16-24 h at 4°C. Samples were subsequently washed with chilled PBS (850 × *g*, 5 min, 4°C), quenched in Quenching Buffer B, and enzymatically dissociated in Dissociation Solution containing collagenase I (0.1 g/10 mL HBSS; 17100-017, Gibco), collagenase IV (0.1 g/10 mL HBSS; LS004188, Worthington), and penicillin-streptomycin (P & S; 208330, Gibco) at 37°C for 40 min with intermittent pipetting every 10 min. The resulting cell suspensions were passed through a 40 μm strainer, washed with PBS, and either processed immediately for library construction or stabilized with Enhancer and 50% glycerol for storage at −80°C. Single-cell libraries were generated using the Chromium Next GEM Single Cell 3′ Gene Expression Flex v2 kit (10x Genomics) and sequenced on an Illumina platform.

### GEM-X Flex single-cell gene expression analysis

Raw sequencing data were processed using CellRanger (v9.0.0, 10x Genomics) and imported into R via the Seurat package (v5). Cells were retained if they contained 300-6,000 detected genes with mitochondrial content below 15%. Following normalization and principal component analysis, batch effects were corrected using Harmony, and cells were clustered at resolution 0.3 and visualized by UMAP. Eleven transcriptionally distinct populations were annotated based on canonical marker gene expression, encompassing luminal and glandular epithelium, stroma, mesothelium, endothelium, pericytes, and immune cells. Differential expressions between *Erbb3^f/f^* and *Erbb3^d/d^*stromal cells were assessed using FindMarkers (logFC > 0.25, min.pct > 0.1), and the resulting gene list was cross-referenced against a curated growth factor gene set (GO:0008083) to identify candidate paracrine factors dysregulated in *Erbb3^d/d^* stroma.

### Re-analysis of published single-cell RNA sequencing data

Raw data were obtained from Wang *et al.* (GSA: CRA011266) ^34^, which profiled mouse uteri at 2.5, 3.5, 4, and 4.5 DPC. In that dataset, 0.5 DPC equals the day of vaginal-plug detection; in our study this corresponds to day 1, thus 2.5 DPC equals day 3 of pregnancy. FASTQ files from 16 sequencing runs (four runs per time point: runs 59-62 for DPC 2.5, 63-66 for DPC 3.5, 67-70 for DPC 4, and 71-74 for DPC 4.5) were processed with Cell Ranger v5.0.1 (10x Genomics) against the mm10 reference genome to generate filtered feature-barcode matrices. Individual Seurat objects were created for each run (Seurat v5.2.1; min.cells = 3, min.features = 200), then merged into a single object with run-specific cell-ID prefixes. Quality control retained cells expressing >500 and <6,000 genes with <15% mitochondrial reads. Data were log-normalized, the top 2,000 variable features were identified by VST, and expression values were scaled. PCA was performed on variable features, followed by Harmony batch correction across runs. UMAP embedding and graph-based clustering (resolution 0.5) were computed using 10 Harmony-corrected dimensions. Clusters were annotated on the basis of established markers into 19 cell types: luminal epithelium (LE), glandular epithelium (GE), stromal subsets (Str_1-Str_4), proliferating stroma (Prol_Str1-Prol_Str3), endothelial (Endo), mesothelial (Meso), pericyte (Peri), myofibroblast (Myo), macrophage subsets (Macro_1-Macro_3), NK cell subsets (NK_1, NK_2), T cells, and B cells. The Peri/Myo cluster was further sub-clustered at resolution 0.1 using Harmony dimensions 1-10 and separated into pericyte (sub-cluster 0; *Notch3^+^/Rgs5^+^*) and myofibroblast (sub-clusters 1 and 2, merged; *Acta2^+^/Tagln^+^*) populations. For day 3-specific analyses, the DPC 2.5 subset was extracted. To focus on the stromal-epithelial signaling axis, stromal subclusters (Str_1-Str_4, Prol_Str1, Prol_Str3) were merged into a single “Str” class, immune populations were merged into “Immune,” and Prol_Str2 was removed. The resulting eight broad cell classes (LE, GE, Str, Meso, Endo, Peri, Myo, Immune) were validated by Spearman correlation heatmap and dot plot of canonical markers.

### CellChat analysis

Cell-cell communication among LE, GE, and Str on day 3 of pregnancy was inferred using CellChat v1.6.1 with CellChatDB.mouse as the ligand-receptor reference. The normalized expression matrix was extracted from the RNA assay data layer, and a CellChat object was created with cell-type labels. Standard workflows were applied: overexpressed genes and interactions were identified, communication probabilities were computed, interactions with fewer than 10 cells in either sender or receiver group were filtered, pathway-level probabilities were calculated, and signaling networks were aggregated. Network centrality metrics were computed via netAnalysis_computeCentrality. Global signaling roles were visualized by scatter plot of outgoing versus incoming strength and by outgoing/incoming signaling pattern heatmaps. Information flow scores for all inferred pathways were ranked by summing interaction probabilities across all sender-receiver pairs per pathway. For the IGF pathway specifically, Str-LE and Str-GE signaling was visualized with circle plots (netVisual_aggregate), IGF-specific outgoing and incoming strengths were quantified by bar plot, receptor expression (*Igf1r, Itga6, Itgb4*) across the three populations was shown by violin plot, and ligand-receptor pair communication probabilities were displayed with a bubble plot. All communication probabilities for inferred interactions were exported as source data.

### Statistical analysis

Statistical analyses were conducted using GraphPad Prism (v8.0) and R (v4.4.1) along with RStudio (2024.04.2). Each experiment was repeated at least three times. Data are shown as mean ± SEM. Statistical analyses were performed using a two-tailed Student’s *t*-test, except for pregnancy outcomes presented in Supplementary Table 1, which were analyzed using a *Chi*-square test. A *P* value less than 0.05 was considered statistically significant.

## Data availability

The raw and processed bulk RNA-seq data generated in this study have been deposited in the NCBI Gene Expression Omnibus (GEO) under accession number GSE309147. The raw and processed single-cell RNA-seq data (generated using the 10x Genomics Chromium Single Cell Gene Expression Flex platform) have been deposited in GEO under accession number GSE309241. Source data are provided with this paper.

## Code availability

The source code for all bioinformatic analyses has been deposited in the Zenodo repository (https://doi.org/10.5281/zenodo.19342033).

## Acknowledgments

We sincerely thank Francesco DeMayo (National Institute of Environmental Health Science, Research Triangle Park, NC) and John B. Lydon (Baylor College of Medicine, Houston, TX) who originally provided *Pgr^Cre/+^*mice, and David Threadgill (Texas A&M University, College Station, TX) for providing *Erbb3* floxed mice. This study was made possible, in part, using the Cincinnati Children’s Single Cell Genomics Facility [RRID:SCR_022653]. We specifically acknowledge the assistance of Kelly Bailie and George Yoshida. Some of the figures in this study were created using BioRender. We thank BioRender.com for providing the platform. This work was supported by NIH grants HD068524 (X.S. and S.K.D.) and HD103475 (S.K.D.). Bo Li is supported by a Lalor foundation postdoctoral fellowship.

## Author contributions

S.K.D. and X.S. designed research; B.L. and A.D. performed research; B.L. and X.S. analyzed data; and B.L., S.K.D., and X.S. wrote the paper.

