## Supplementary Information for "Uterine ERBB3 signaling is critical to cultivate stromal environment for functional gland branching"

**This PDF file includes:**

Supplementary Tables 1-2

Supplementary Figs. 1-12

**Supplementary Table 1:**

**Pregnancy outcomes in *Erbb3*$^{f/f}$, *Erbb3*$^{f/f}$ *Ltf*$^{Cre/+}$, and *Erbb3*$^{f/f}$ *Pgr*$^{Cre/+}$ mice.**

| | No. of plug positive mice | No. of mice with pups | Total No. of pups | No. of pups/litter (mean ± SEM) | No. of mice with pups/ No. of plug positive mice (%) |
|---|---|---|---|---|---|
| *Erbb3*$^{f/f}$ | 12 | 10 | 59 | 5.9 ± 0.4 | 83.3 |
| *Erbb3*$^{f/f}$ *Ltf*$^{Cre/+}$ | 10 | 9 | 54 | 6.0 ± 0.3 | 90 |
| *Erbb3*$^{f/f}$ *Pgr*$^{Cre/+}$ | 10 | 3 | 15 | 5.0 ± 1.0 | 30[a] |

[a]$p < 0.05$ when compared to *Erbb3*$^{f/f}$ mice; chi-squared test.

**Supplementary Table 2:**

**Sequences of primers used for qPCR analyses.**

| Genes | Sequence (5' to 3') |
|---|---|
| *Erbb3-Mus-F* | *GTGCTGGGTTTCCTTCTCAG* |
| *Erbb3-Mus-R* | *ACCCATGACCACCTCACACT* |
| *Gapdh-Mus-F* | *AGGTCGGTGTGAACGGATTTG* |
| *Gapdh-Mus-R* | *TGTAGACCATGTAGTTGAGGTCA* |
| *Lif-Mus-F* | *AAAGCTATGTGCGCCTAA* |
| *Lif-Mus-R* | *ACCATCCGATACAGCTCCAC* |

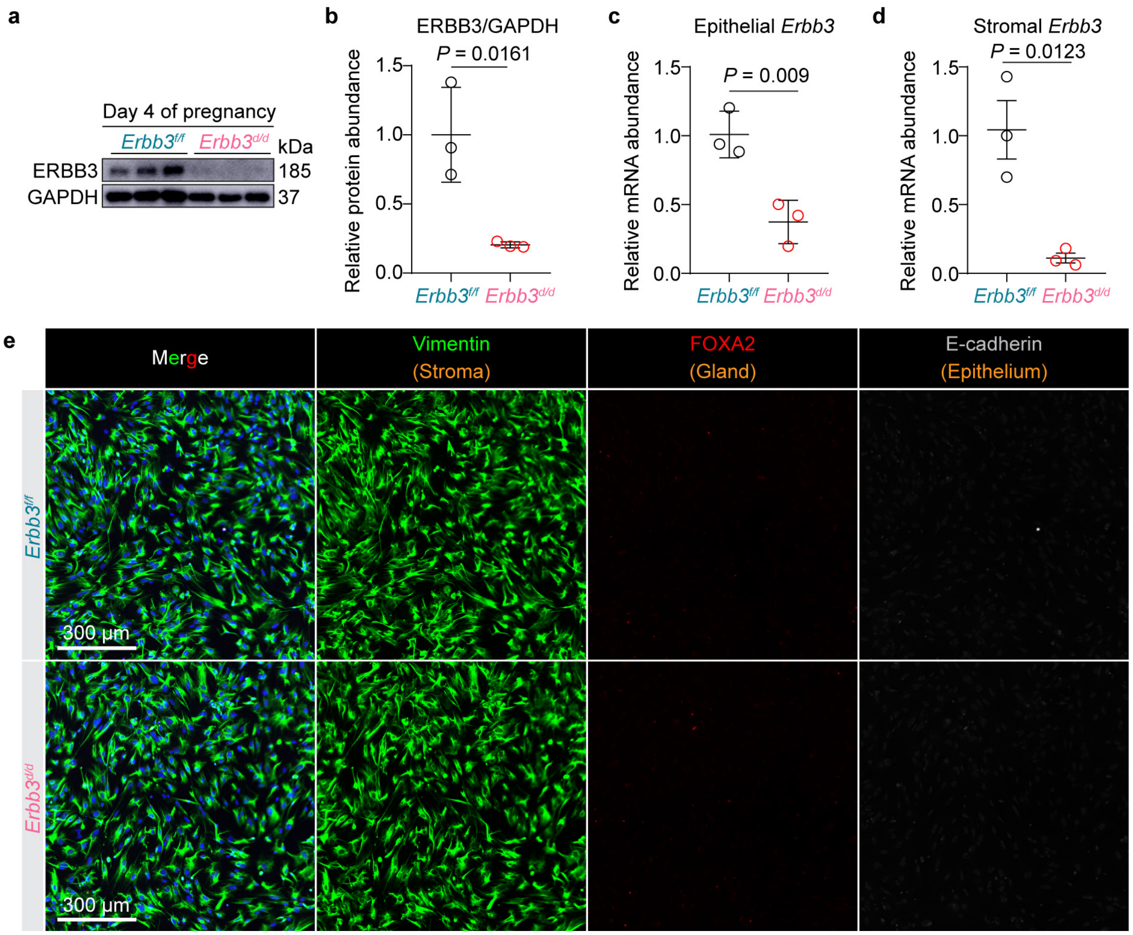


**Supplementary Fig. 1. Efficient deletion of *Erbb3* and purity validation in isolated uterine cells on day 4 of pregnancy.** **a** Western blot analysis of ERBB3 in day 4 pregnant uteri from *Erbb3*$^{f/f}$ (n = 3) and *Erbb3*$^{d/d}$ (n = 3) mice. GAPDH served as a loading control. **b** Quantitative analysis of relative ERBB3/GAPDH protein abundance. Data are represented as mean ± SEM. *P* values were determined by a two-tailed Student's *t* test. Source data are provided as a Source Data file. **c, d** qPCR analysis of *Erbb3* mRNA abundance in isolated primary uterine epithelial (**c**) and stromal (**d**) cells from *Erbb3*$^{f/f}$ (n = 3) and *Erbb3*$^{d/d}$ (n = 3) mice on day 4 of pregnancy. Data are represented as mean ± SEM. *P* values were determined by a two-tailed Student's *t* test. Source data are provided as a Source Data file. **e** Immunofluorescence characterization of the isolated stromal cell fractions to confirm lineage purity. Cultured cells from both genotypes were co-stained for Vimentin (stromal marker), FOXA2 (glandular epithelial marker), and E-cadherin (pan-epithelial marker). Nuclei were counterstained with DAPI. Scale bars, 300 μm. Representative images from n = 3 independent biological replicates per group are shown.

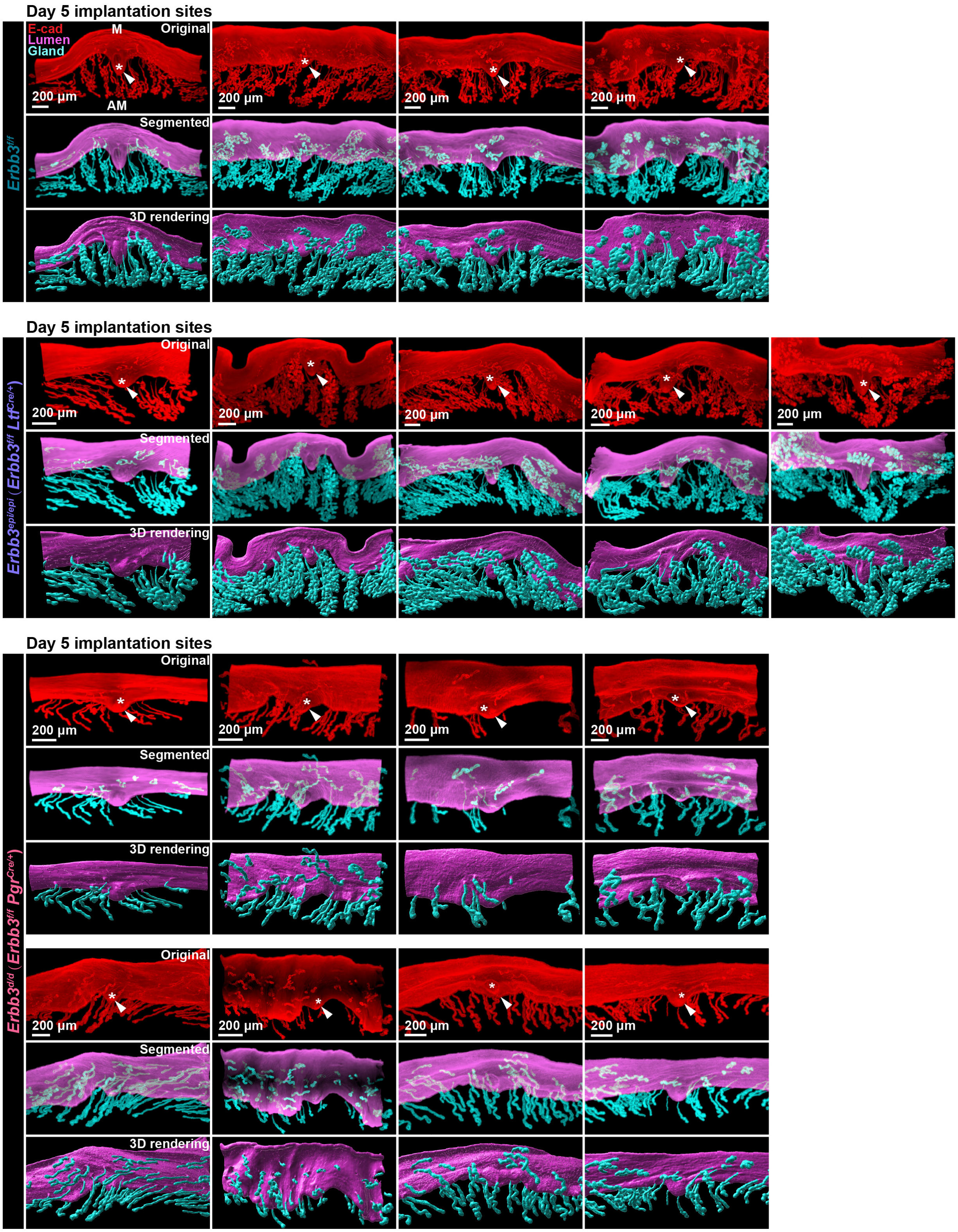


**Supplementary Fig. 2. 3D images of day 5 implantation sites from *$Erbb3^{f/f}$*, *$Erbb3^{epi/epi}$*, and *$Erbb3^{d/d}$* mice.** 3D imaging of day 5 implantation sites in *$Erbb3^{f/f}$* (n = 4), *$Erbb3^{epi/epi}$* (n = 5), and *$Erbb3^{d/d}$* (n = 8) females. Images of E-cadherin immunostaining, segmented, and 3D rendering of day 5 implantation sites in each genotype. Scale bars, 200 µm. Asterisks indicate the location of blastocysts. Arrowheads indicate the implantation chamber (crypt). M mesometrial pole, AM antimesometrial pole.

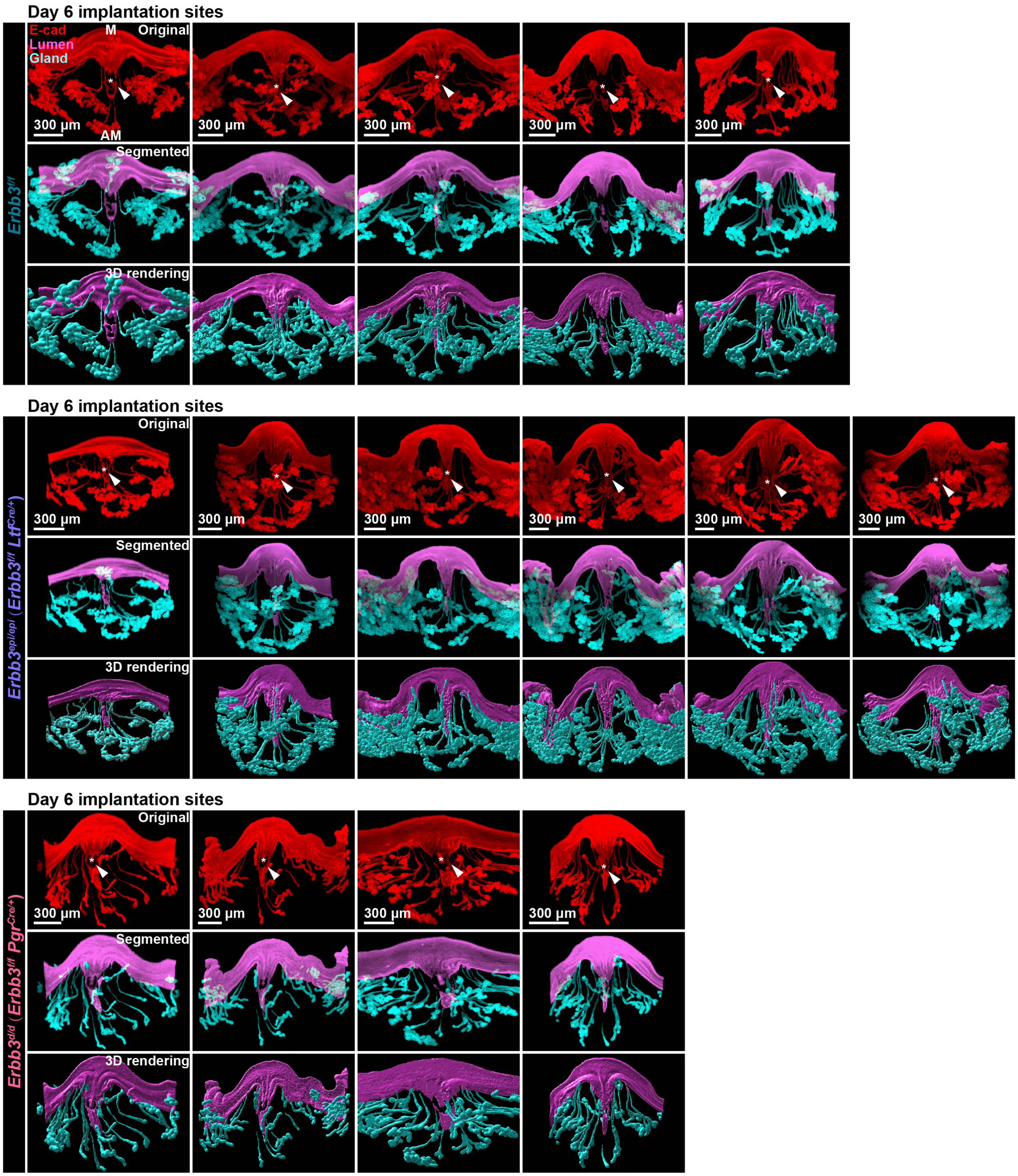


**Supplementary Fig. 3. 3D images of day 6 implantation sites from *Erbb3*$^{f/f}$, *Erbb3*$^{epi/epi}$, and *Erbb3*$^{d/d}$ mice.** 3D imaging of day 6 implantation sites in *Erbb3*$^{f/f}$ (n = 5), *Erbb3*$^{epi/epi}$ (n = 6), and *Erbb3*$^{d/d}$ (n = 4) females. Images of E-cadherin immunostaining, segmented, and 3D rendering of day 6 implantation sites in each genotype. Scale bars, 300 µm. Asterisks indicate the location of blastocysts. Arrowheads indicate the implantation chamber (crypt). M mesometrial pole, AM antimesometrial pole.

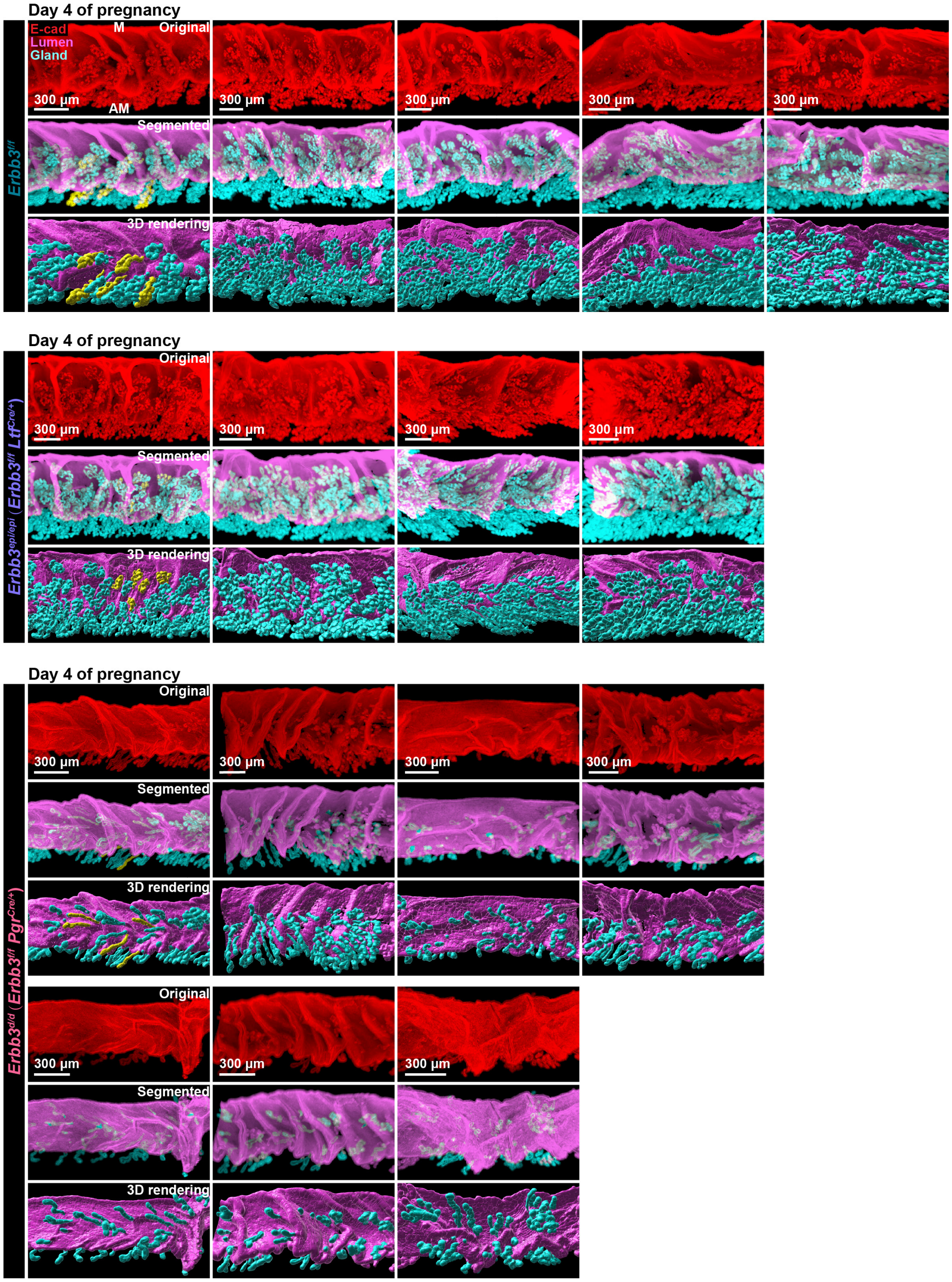


**Supplementary Fig. 4. 3D images of day 4 mouse uteri from *Erbb3*$^{f/f}$, *Erbb3*$^{epi/epi}$, and *Erbb3*$^{d/d}$ mice.** 3D imaging of day 4 mouse uteri in *Erbb3*$^{f/f}$, *Erbb3*$^{epi/epi}$, and *Erbb3*$^{d/d}$ females. Images of E-cadherin immunostaining, segmented, and 3D rendering of day 4 mouse uteri in each genotype. Scale bars, 300 µm. M, mesometrial pole; AM, antimesometrial pole.

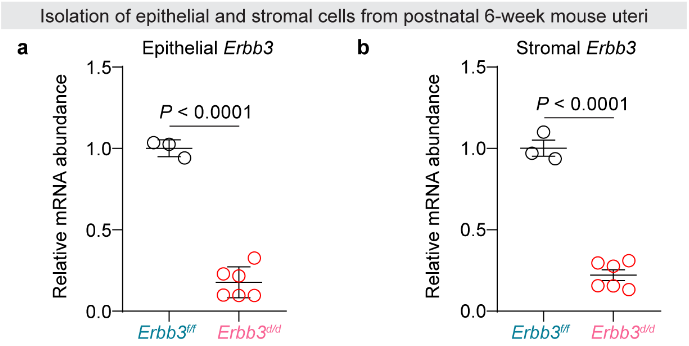


**Supplementary Fig. 5. Efficient deletion of *Erbb3* in isolated uterine cells at postnatal 6 weeks of age. a, b** qPCR analysis of *Erbb3* mRNA abundance in isolated primary uterine epithelial (**a**) and stromal (**b**) cells from *Erbb3*$^{f/f}$ (n = 3) and *Erbb3*$^{d/d}$ (n = 6) mice at postnatal 6 weeks of age. Data are represented as mean ± SEM. *P* values were determined by a two-tailed Student's *t* test. Source data are provided as a Source Data file.

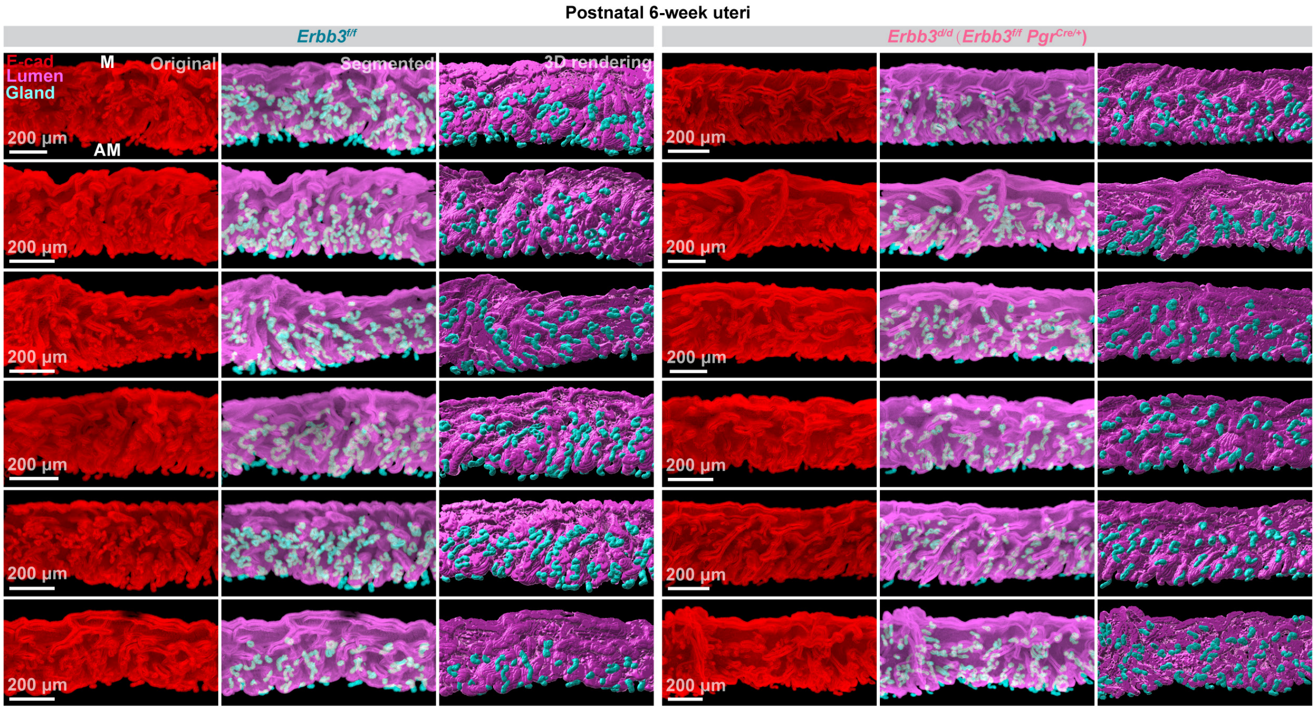


**Supplementary Fig. 6. 3D images of postnatal 6-week mouse uteri from *Erbb3*$^{f/f}$ and *Erbb3*$^{d/d}$ mice.** 3D imaging of postnatal 6-week mouse uteri in *Erbb3*$^{f/f}$ (n = 6) and *Erbb3*$^{d/d}$ (n = 6) females. Images of E-cadherin immunostaining, segmented, and 3D rendering of 6-week mouse uteri in each genotype. Scale bars, 200 µm. M, mesometrial pole; AM, antimesometrial pole.

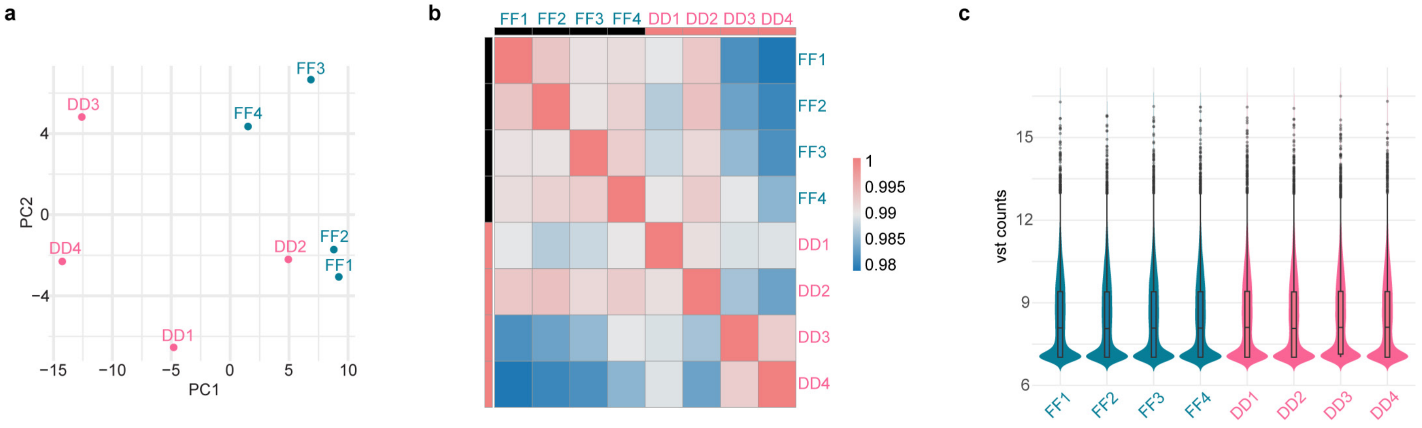


**Supplementary Fig. 7. Principal component analysis (PCA) of uterine stromal bulk RNA-seq samples from *Erbb3*$^{f/f}$ and *Erbb3*$^{d/d}$ mice. a** PCA plot showing transcriptomic variation among stromal samples isolated from *Erbb3*$^{f/f}$ (n = 4) and *Erbb3*$^{d/d}$ (n = 4) uteri on day 3 of pregnancy. The first two principal components (PC1 and PC2) are plotted, revealing clear separation between genotypes. **b** Sample to sample correlation heatmap based on variance-stabilizing transformed (vst) counts, showing high within-group similarity and clustering by genotype. **c** Violin plots of vst-transformed gene expression distributions across stromal RNA-seq samples, indicating comparable overall expression profiles across samples.

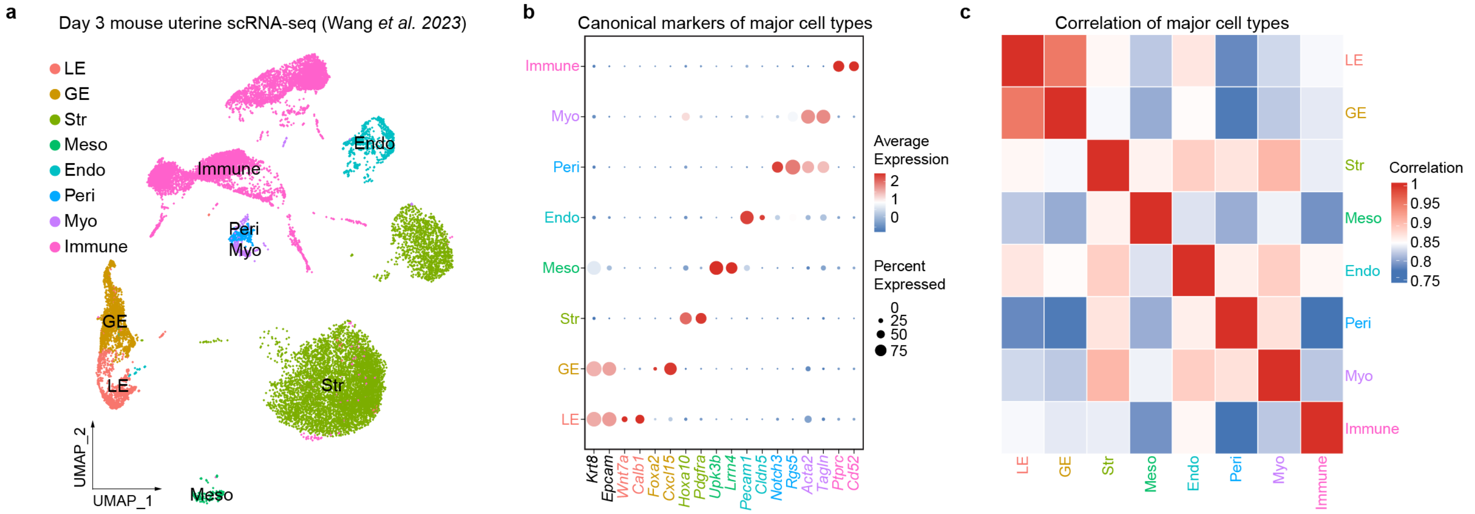


**Supplementary Fig. 8. Single-cell transcriptomic reanalysis identifies distinct uterine cell populations on day 3 of pregnancy. a** UMAP visualization of reanalyzed day 3 pregnant mouse uterine cells, identifying major distinct compartments including luminal epithelium (LE), glandular epithelium (GE), and stroma (Str). **b** Dot plot of canonical marker genes validating the annotated cell clusters. Dot size indicates the percentage of expressing cells; color intensity represents average expression. **c** Correlation heatmap demonstrates overall transcriptional similarities among the identified cell types.

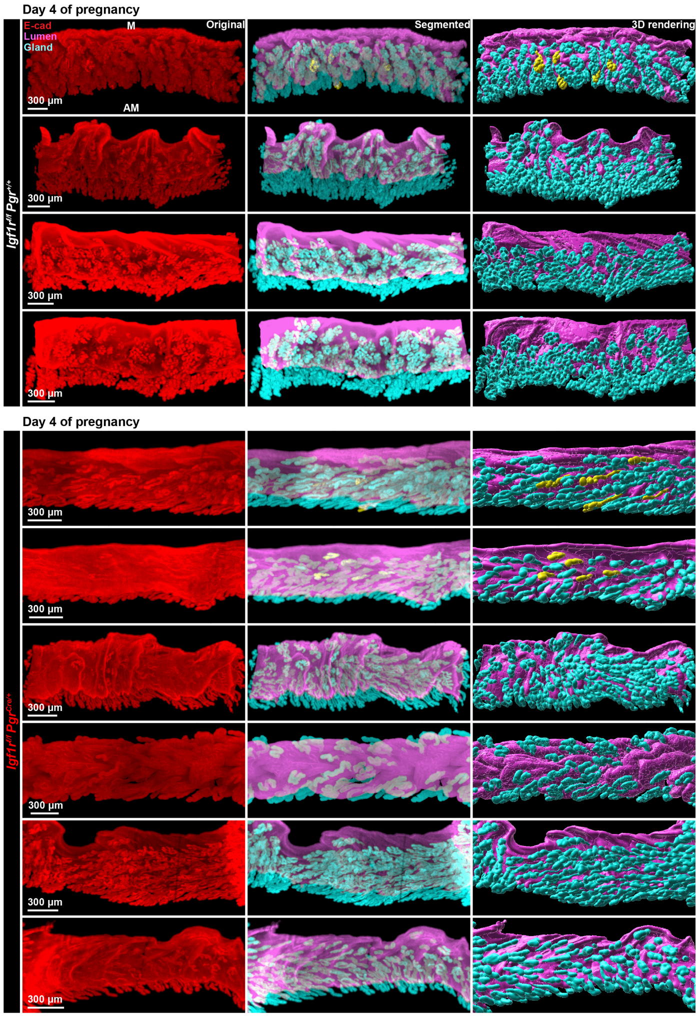


**Supplementary Fig. 9. *Igf1r* deficiency impairs uterine gland branching.** Whole-mount 3D imaging of day 4 pregnant uteri from *Igf1r*$^{f/f}$ *Pgr*$^{+/+}$ (n = 4) and *Igf1r*$^{f/f}$ *Pgr*$^{Cre/+}$ (n = 6) females. From left to right, columns display original E-cadherin immunostaining, segmented (lumen in magenta, glands in cyan), and 3D surface rendering. M, mesometrial pole; AM, antimesometrial pole. Scale bars: 300 μm.

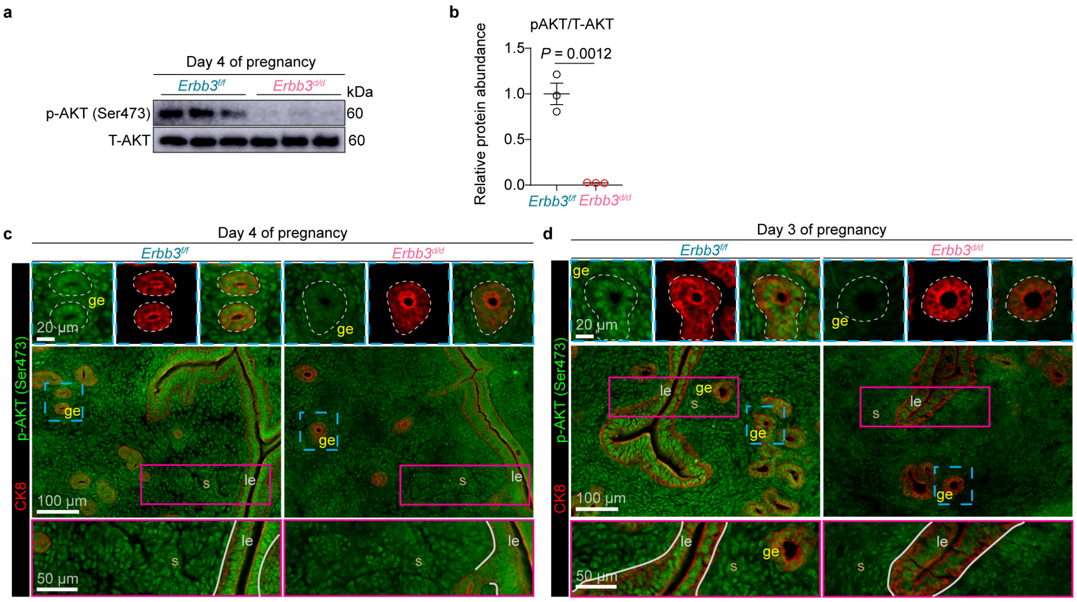

**Supplementary Fig. 10. Uterine deletion of *Erbb3* suppresses AKT signaling in the *Erbb3*$^{d/d}$ uteri. a** Western blot analysis of phosphorylated AKT (p-AKT, Ser473) and total AKT (T-AKT) in day 4 pregnant uteri from *Erbb3*$^{f/f}$ (n = 3) and *Erbb3*$^{d/d}$ (n = 3) mice. GAPDH served as a loading control. **b** Quantitative analysis of relative p-AKT/T-AKT (**a**) protein abundance. Data are represented as mean ± SEM. *P* values were determined by a two-tailed Student's *t* test. Source data are provided as a Source Data file. **c, d** Representative immunofluorescence staining for p-AKT (Ser473) and the epithelial marker CK8 in *Erbb3*$^{f/f}$ and *Erbb3*$^{d/d}$ uteri on day 4 (**c**) and day 3 (**d**) of pregnancy (n = 3 mice per group per time point). Top insets highlight the glandular epithelium (ge, outlined by yellow dashed lines). Bottom insets detail the stroma (s) and luminal epithelium (le). Scale bars: 100 μm (main images), 50 μm (bottom insets), and 20 μm (top insets).

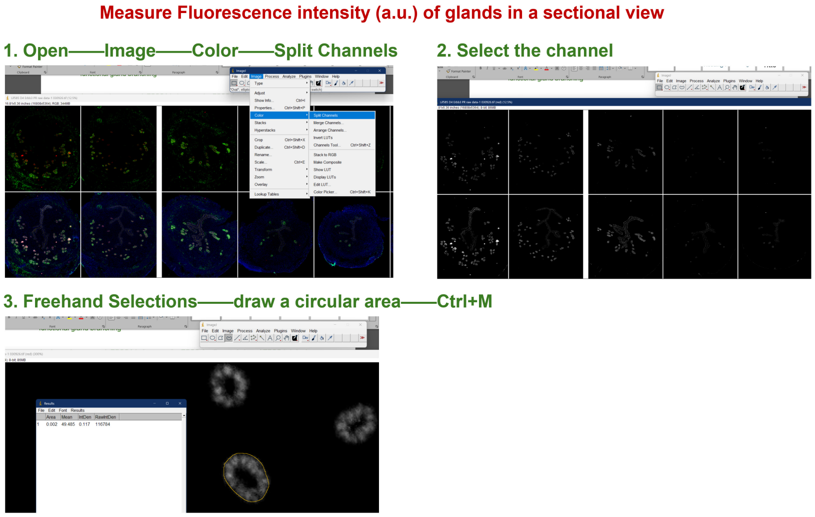


**Supplementary Fig. 11. Workflow for quantification of fluorescence intensity in uterine tissues.** Representative images illustrating the ImageJ-based procedure used to measure fluorescence intensity (arbitrary units, a.u.) of uterine tissues in sectional view.

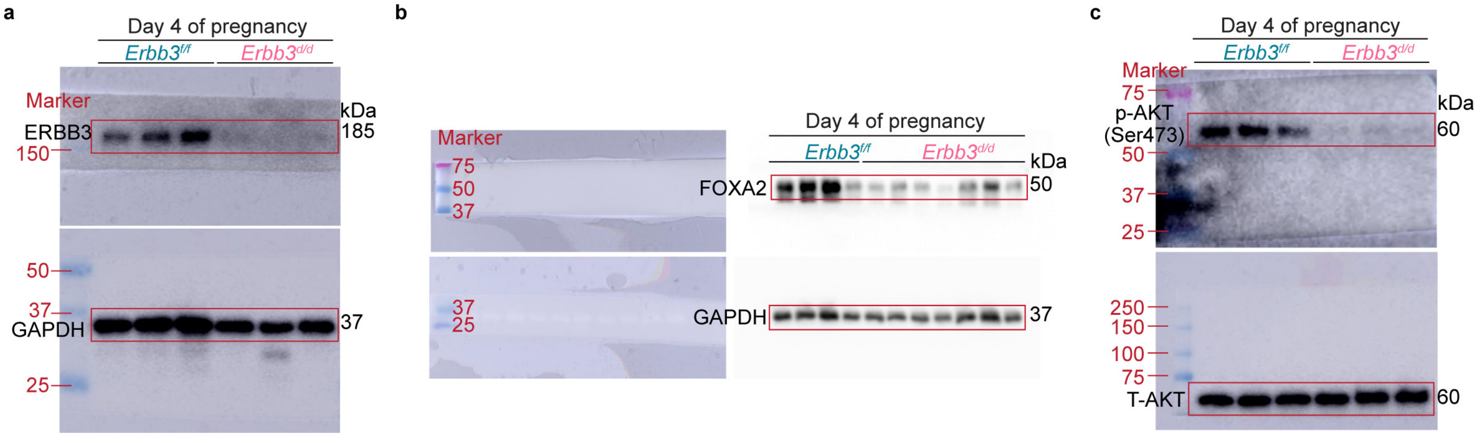


**Supplementary Fig. 12. Uncropped western blot images for all blots presented in this study. a** Western blot image corresponding to Supplementary Fig. 1a. **b** Western blot images corresponding to Fig. 2k. **c** Western blot image corresponding to Supplementary Fig. 10a.
